# Distinct synchronization, cortical coupling and behavioural function of two basal forebrain cholinergic neuron types

**DOI:** 10.1101/703090

**Authors:** Tamás Laszlovszky, Dániel Schlingloff, Panna Hegedüs, Tamás F. Freund, Attila Gulyás, Adam Kepecs, Balázs Hangya

## Abstract

Basal forebrain cholinergic neurons (BFCN) densely innervate the forebrain and modulate synaptic plasticity, cortical processing, brain states and oscillations. However, little is known about the functional diversity of cholinergic neurons and whether distinct types support different functions. To examine this question we recorded BFCN *in vivo*, to examine their behavioral functions, and *in vitro*, to study their intrinsic properties. We identified two distinct types of BFCN that markedly differ in their firing modes, synchronization properties and behavioral correlates. Bursting cholinergic neurons (BFCN_BURST_) fired in zero-lag synchrony with each other, phase-locked to cortical theta activity and fired precisely timed bursts of action potentials after reward and punishment. Regular firing cholinergic neurons (BFCN_REG_) were found predominantly in the posterior basal forebrain, displayed strong theta rhythmicity (5-10 Hz), fired asynchronously with each other and responded with precise single spikes after behavioral outcomes. In an auditory detection task, synchronization of BFCN_BURST_ neurons to auditory cortex predicted the timing of mouse responses, whereas tone-evoked cortical coupling of BFCN_REG_ predicted correct detections. We propose that cortical activation relevant for behavior is controlled by the balance of two cholinergic cell types, where the precise proportion of the strongly activating BFCN_BURST_ follows an anatomical gradient along the antero-posterior axis of the basal forebrain.

## Introduction

Basal forebrain cholinergic neurons have been associated with learning, memory, plasticity, attention, arousal, regulation of food intake, sleep-wake cycle and even consciousness^1–10^. These processes are modulated at different time scales; indeed, the cholinergic system was hypothesized to exhibit both slow tonic and fast phasic effects^11–13^, similar to the dopaminergic or noradrenergic systems^14,15^. *In vitro* studies associated heterogeneous firing patterns with varying temporal scales, suggesting that subtypes of cholinergic neurons may underlie this temporal and functional heterogeneity, i.e. early firing cholinergic neurons are dedicated to phasic activation and late firing neurons fire slowly in order to set ambient acetylcholine levels^16–21^. However, *in vivo* studies^22–24^ have not examined the functional heterogeneity of cholinergic neurons. Therefore, we tested whether there are distinct types of BFCN *in vivo* and *in vitro*, and if so, whether these types show differences in responding to behaviorally salient events, synchronizing within and across types as well as with cortical activities.

Inter-areal synchrony has been proposed as a hallmark of neural communication and efficient information transfer. Distant brain areas can engage in synchronous oscillations, and this oscillatory activity is thought to orchestrate neuronal processing^25–28^. This clock-like organization results in the phase locking of neuronal spiking to ongoing oscillations at the cellular level, different patterns of synchrony within and across cell types at the network level, and rhythmic fluctuation of sensory detection^29–32^, attention^33–37^, and reaction time^33,38^ at the behavioral level. Therefore, synchronous versus asynchronous activation of subcortical inputs may have substantially different impact on cortical functions including plasticity, attention, learning and other aspects of cognition. However, recording of pairs of cholinergic neurons simultaneously has not been carried out and thus synchrony between individual cholinergic units has not yet been tested. In addition, assessment of synchrony between cholinergic firing and cortical oscillations has been sparse and seemingly contradictory^39–41^.

We found two distinct cholinergic cell types in the basal forebrain by recording BFCN both *in vivo* and *in vitro*. Bursting cholinergic neurons (BFCN_BURST_) exhibited early firing in response to current injections *in vitro*, strongly bursting or irregular ‘Poisson-like’ firing *in vivo*, within-cell type synchrony and strong correlation to cortical theta oscillation. Characteristically, these neurons fired rapid, brief bursts of action potentials after reward and punishment in an auditory detection task. Synchrony between BFCN_BURST_ and auditory local field potentials (LFP) predicted mouse response but did not differentiate correct and erroneous responses. In addition, we uncovered a unique cholinergic, regular rhythmic type (BFCN_REG_) in the posterior basal forebrain. This type showed late firing in response to current injections *in vitro* that could not be transformed into burst mode. BFCN_REG_ showed largely asynchronous firing with other BFCN. In contrast to BFCN_BURST_, synchronization of BFCN_REG_ with auditory cortical LFP specifically predicted correct responses of mice. Therefore, these differences in firing mode, synchrony and anatomical distribution might underlie the differential regulation of behavior by distinct cholinergic cell types.

## Results

### Distinct firing patterns of cholinergic neurons in vivo

We performed extracellular tetrode recordings from the basal forebrain (BF) of awake mice (Fig.1a)^42^. Cholinergic neurons were identified using an optogenetic tagging approach. Neurons responding with statistically significant short latency firing (Stimulus-Associated spike Latency Test, SALT; p < 0.01) to blue laser light in transgenic mice expressing the photosensitive channelrhodopsin (ChAT-Cre infected by AAV-DIO-EF1a-ChETA, N = 15, by AAV-DIO-EF1a-hChR2(H134R), N = 3 or ChAT-ChR2, N = 3 mice) were considered optogenetically identified cholinergic neurons (n = 56). In addition, neurons that fell in the same cluster by hierarchical clustering of response properties^24,43,44^ were considered putative cholinergic neurons (n = 22; the algorithm was detailed in ref. ^24^). We detected no systematic differences between optogenetically identified and putative cholinergic neurons (Fig.S1, see also Fig.S4 in ref. ^24^), therefore these neurons were pooled and resulted in a data set of 78 BF cholinergic neurons.

**Figure 1.**
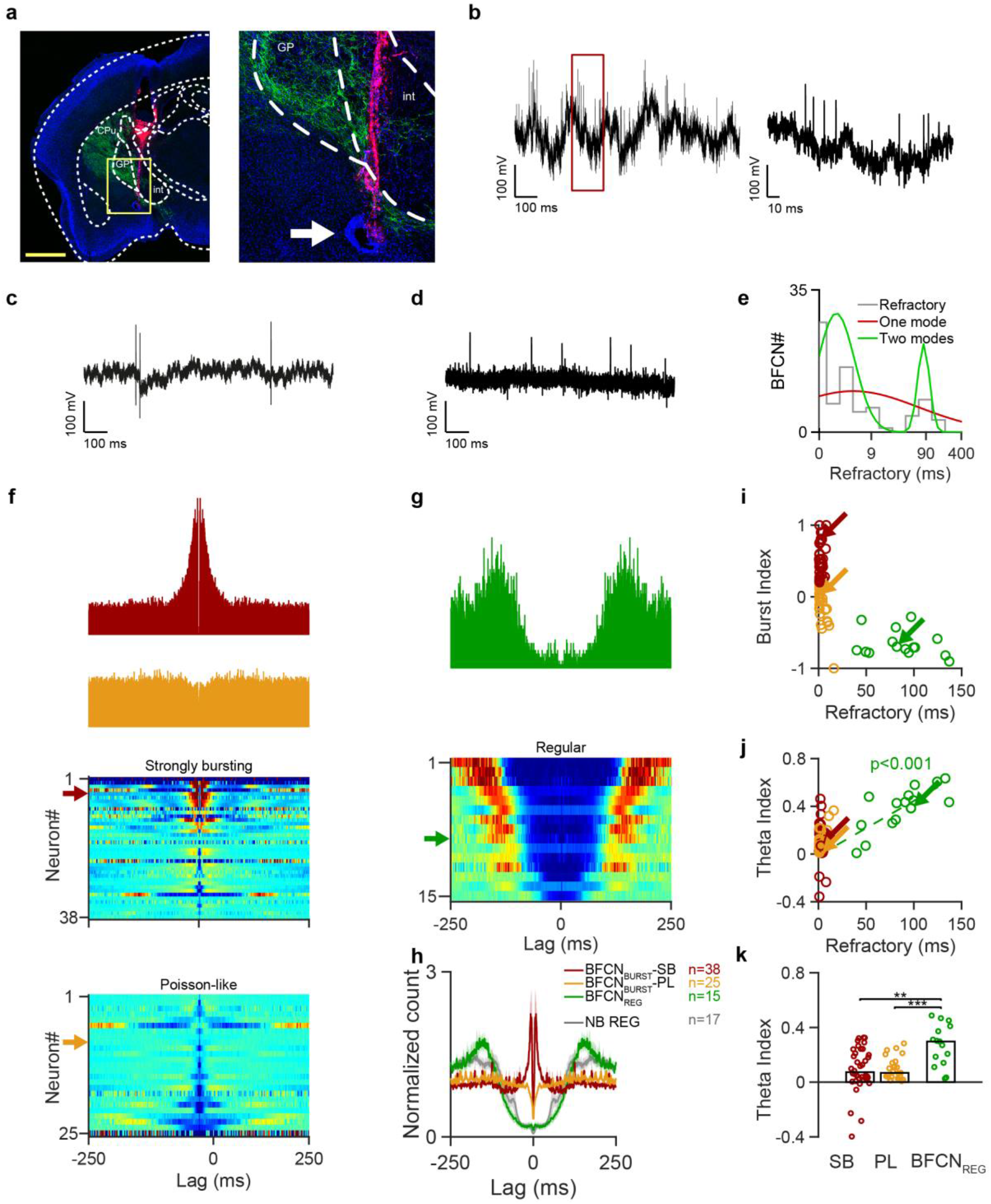
*In vivo* recordings revealed two types of central cholinergic neurons, BFCN_BURST_ and BFCN_REG_. **a,** Coronal section with the tetrode tracks (red, DiI) and the electrolytic lesion (arrow). Scale bar, 1 mm. Green, ChAT-ChR2-Eyfp; blue, nuclear staining (DAPI). **b,** Left, example raw trace of a BFCN_BURST_. Right, burst enlarged. **c,** Example short inter-spike interval of a BFCN_BURST_-PL. **d,** Example raw trace of a BFCN_REG_. **e,** Distribution of relative refractory periods. Log of the refractories (grey bars) were fitted with one (red) or two modes (green). **f,** Top, example auto-correlations of BFCN_BURST_-SB (red) and BFCN_BURST_-PL (orange). Bottom, all neurons as individual rows. **g,** Top, example auto-correlation of a BFCN_REG_ (green); bottom, all neurons as individual rows. **h,** Average auto-correlations. Unidentified regular firing neurons (black) are few and resemble BFCN_REG_ (green) based on their auto-correlograms. **i,** Scatter plot showing Burst Index and relative refractory period. Arrows indicate example cells in f and g. **j,** Correlation between relative refractory period and Theta Index (p = 0.0008 for BFCN_REG_). **k,** Median Theta Index.

Previous *in vitro* studies suggested that cholinergic neurons may exhibit heterogeneous firing patterns^16,19,20^; however, this has not been tested *in vivo* and the potential diversity of BFCN is unexplored in awake animals. We noticed that some cholinergic neurons were capable of firing bursts of action potentials *in vivo* with short, <10 ms inter-spike intervals (ISI), while others exhibited a markedly different pattern of regular rhythmic firing dominated by long inter-spike intervals (Fig.1b-d). To quantify this, we defined relative refractory periods of BF cholinergic neurons based on their auto-correlograms, characterized by low probability of firing (inspired by Royer et al., see Methods^45^). The distribution of relative refractory period duration covered a broad range (1-137 ms) and showed a bimodal distribution with two distinct, approximately log-normal modes^46^ (Fig.1e-h). This was confirmed by a model selection approach based on Akaike and Bayesian information criteria (Fig.S1i). This demonstrated the existence of a separate short-refractory, burst firing and long-refractory, regular firing group of cholinergic neurons. Therefore we coined these cholinergic neurons BFCN_BURST_ and BFCN_REG_, respectively.

We further analyzed the burst firing properties of BFCN_BURST_ and found considerable heterogeneity based on their spike auto-correlations. Many short-refractory neurons exhibited strong bursting patterns with classical ‘burst shoulders’^45^ in their auto-correlograms (BFCN_BURST_-SB, strong bursting), while others showed irregular patterns of inter-spike intervals, resembling a Poisson process (BFCN_BURST_-PL, ‘Poisson-like’; Fig.1f). Of note, the lack of a central peak in the auto-correlation did not preclude the occasional presence of bursts (Fig.1c). These firing patterns were distinct on average (Fig.1h); however, this separation was less evident than the bimodal relative refractory distribution and a few neurons could have been categorized in either group (Fig. 1i).

Interestingly, the long-refractory neurons exhibited strong rhythmicity in the theta frequency band (5-10 Hz; Fig. 1j-k). The strength of rhythmic firing, quantified based on auto-correlation peaks in the theta band (Theta Index, see Methods), was correlated with the length of the relative refractory period (p = 0.0008).

Next, we analyzed the firing patterns of a large dataset of un-tagged BF neurons. Burst firing has been shown for GABAergic BF neurons before^41,47^; in accordance, we found that many non-cholinergic cells were capable of burst firing (Fig.S2a-b). Surprisingly however, only a small proportion of untagged BF neurons showed regular rhythmic firing with long refractory period (n = 17; Fig.S2c-g), which were similar to those that we had characterized as cholinergic (n = 12; Fig.1h). This suggests that at least about 40% of regular rhythmic BF neurons are cholinergic and may provide means to identify this subgroup of putative cholinergic neurons based on firing rate and regular rhythmic activity pattern when response to air puff is not available (Fig. S2h).

### In vitro recordings confirmed two types of cholinergic neurons

We were wondering whether the cholinergic firing patterns uncovered by our *in vivo* recordings reflect intrinsic properties, hence these neurons can be thought of as distinct cell types. Alternatively, distinct firing patterns may be determined by the current state of the network or variations in the input strength of individual cells. To answer this we turned to *in vitro* preparations, where the membrane potential of the neuron and the strength of activation are precisely controlled and monitored.

We performed whole cell patch clamp recordings from n = 60 cholinergic neurons from the basal forebrain in acute slices. Cholinergic neurons were identified by their red epifluorescence in N = 12 mice injected with AAV2/5-EF1a-DIO-hChR2(H134R)-mCherry-WPRE-HGHpA (Fig.2a). We applied a somatic current injection protocol (Fig.2b) containing a 3-second-long incremental ‘prepolarization’ step followed by a positive square pulse (1 s), to elicit spiking starting from different membrane potentials. We found two distinct behaviors upon current injection (Fig.2b-i) using similar testing conditions (Fig.S3a). Cholinergic cells from the first group (red, n = 29) displayed short spike delay (8.05 ± 0.74 ms, median ± SE of median) and bimodal ISI distribution with short ISIs corresponding to high-frequency ‘burst’ firing (maximum, 122.69 ± 18.99 Hz; Fig.2h-i). The second group (green, n = 31) displayed low maximal firing rate (13.81 ± 2.32 Hz, p < 0.0001), unimodal ISI histogram, and a prominent spike delay (maximum spike delay, 153.05 ± 55.59 ms, p < 0.0001 compared to first group) which depended on the membrane potential prior to spiking (Fig.2f-g). Importantly, depolarized late-firing cells responded to suprathreshold current injections with short spike delay opposed to hyperpolarized state where late firing was prominent (Fig.S3b). These distinct early responding / burst firing or late responding / non-bursting modes were also reliably elicited by optogenetic depolarization (Fig.2c-d). Spontaneous action potentials revealed shorter spikes and large amplitude slowly decaying AHP in late-compared to early firing (bursting) cells (Fig.2e). To compare *in vivo* and *in vitro* firing patterns, we calculated auto-correlations and Burst Indices (early firing, 0.64 ± 0.08; late firing, −1.0 ± 0, p < 0.0001) from spike trains during the current injection protocol (Fig. 2j-k). Early and late firing neurons *in vitro* matched BFCN_BURST_ and BFCN_REG_ *in vivo*, suggesting these groups are the same.

**Figure 2.**
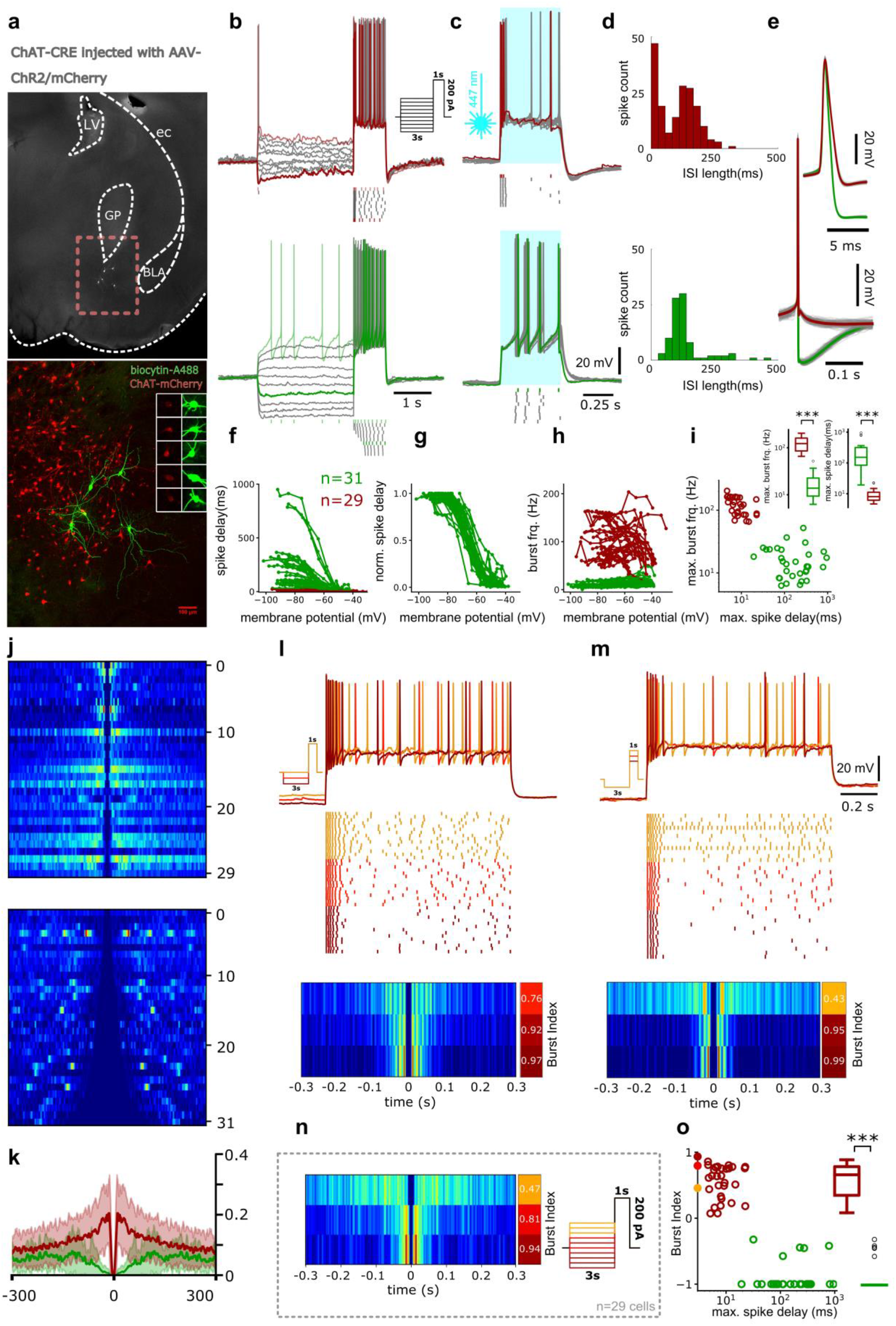
*In vitro* recordings confirmed two types of central cholinergic neurons. **a,** Representative confocal image of recorded and biocytin-filled cholinergic cells expressing ChR2 in nucleus basalis of the BF. **b,** Top, firing pattern of an early firing cell showing short spike delay and high-frequency spike clusters upon positive current injections. Bottom, firing pattern of a representative late-firing cholinergic cell showing low maximal firing rate and prominent spike delay when driven from hyperpolarized membrane states. **c,** The same cells show similar responses upon photostimulation (0.5 s). **d,** Inter-spike interval histograms of the same cells show bimodal (early firing) or unimodal (late firing) distributions. **e,** Average action potential shape from an example early- (red) and late-firing (green) cholinergic cell (100 APs/cell in grey, average in color). **f,** Spike delay depended on membrane potential. **g,** Normalized spike delay showed stereotypic behavior in late firing cholinergic neurons. **h,** Maximal firing rate as a function of membrane potential. **i,** Maximal firing rate plotted against maximal spike delay in all recorded cells. **j,** Spike auto-correlograms during somatic current injection protocols. **k,** Averaged auto-correlograms of early- (red) and late-firing (green) cholinergic cells (solid line, mean; shading, SEM). **l,** Firing pattern of an early-firing cell in response to three current injection protocols with different current magnitude applied prior to depolarization step. The protocol was designed to model internal state (membrane potential) dependence of spiking pattern in response to uniform input. Raster plot represents 20 trials with each protocol (deeper red, more hyperpolarized states). Bottom, corresponding auto-correlograms and Burst Indices. **m,** Firing pattern of the same cell in response to current injection protocols with different depolarization step magnitude. The protocol was designed to mimic input strength dependence of spiking pattern. Raster plot shows 20 trials for each protocol with deeper red corresponding to smaller injected currents. **n,** Average auto-correlations and corresponding Burst Indices of early-firing cells (n=29) driven from depolarized (top) and hyperpolarized (bottom) states. Three groups were formed from all early firing cells based on the 3-second-long ‘pre-polarization’ magnitude (right inset). **o,** Burst Index plotted against maximal spike delay. Dots overlaid on y-axis correspond to Burst Indices presented on panel n.

Next, we tested whether the different *in vivo* firing modes of bursting cholinergic neurons (BFCN_BURST_-SB vs. BFCN_BURST_-PL) could be explained by variations in membrane potential and input strength. To investigate this possibility we applied somatic current injection protocols designed to test input and state dependency of burstiness. Indeed, we found that the same BFCN_BURST_ cells were capable of producing both strongly bursting and Poisson-like firing patterns. This property depended both on the membrane potential of the neuron (Fig.2l,n-o) and the strength of the activation (Fig.2m), with Poisson-like firing occurring more frequently at more depolarized states and in response to stronger depolarizing inputs. In summary, we identified two types of BFCN. BFCN_REG_ showed regular theta-rhythmic firing *in vivo* and late, regular responses to current injections *in vitro*; BFCN_BURST_ exhibited burst firing both *in vivo* and *in vitro*, where the strength of bursting was determined by the level of excitation.

### Cholinergic bursts transmit phasic information about reinforcers

Cholinergic neurons act at different timescales regulating different aspects of cognition from slow sleep-wake and arousal processes to fast subsecond or even millisecond timescales of reinforcement learning and plasticity^4,12,13,24,48^. Based on *in vitro* studies it was hypothesized that bursting specifically represents fast ‘phasic’ information transfer^16^; however, this has not been tested. Therefore, we analyzed the activity of BF cholinergic neurons after reward and punishment in mice performing auditory conditioning^24^ (Fig.3a).

**Figure 3.**
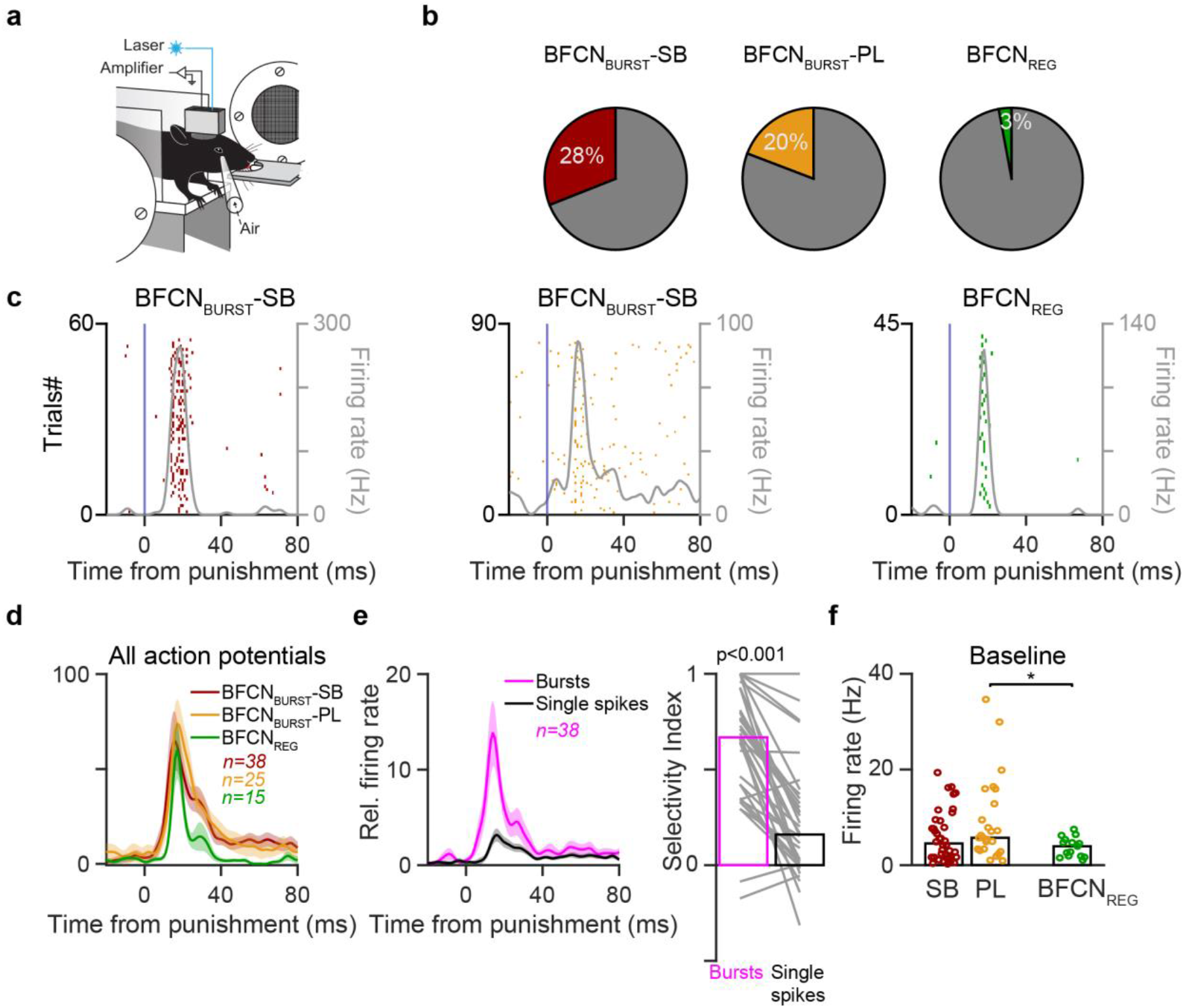
Cholinergic bursts transmit phasic information about reinforcers. **a,** Mice were trained to lick for cue stimuli of pure tones. Hits were rewarded with a drop of water, whereas False Alarms were punished by air puff. Modified from ref. ^24^ **b,** Percentage of intra-burst action potentials. **c,** Example raster plots of phasic responses to punishment by BFCN_BURST_ (left) and BFCN_REG_ (right). Peri-event time histograms smoothed by moving average are overlaid in grey. **d,** Average response of cholinergic neurons to punishment. **e,** Left, occurrence of bursts and single spikes in BFCN_BURST_-SB normalized to baseline; right, Selectivity Index calculated as spike number in 20-50 relative to 100-250 ms post-event windows. Bursts of BFCN_BURST_-SB are more concentrated after punishment compared to single spikes. **f,** Median baseline firing rate.

We defined a burst as a series of action potentials starting with an ISI below 10 ms and subsequent ISIs below 15 ms to allow for typical ISI accommodation patterns^45^. As expected, BFCN_BURST_ categorized based on auto-correlograms showed a high percentage of burst firing: 28% for BFCN_BURST_-SB and 20% for BFCN_BURST_-PL, while little burst activity was detected in the BFCN_REG_ (3%, Fig. 3b).

We have shown previously that the strongest response of cholinergic neurons occurred after air puff punishment^24^: BFCN responded phasically with short latency (18 ± 1.9 ms, median ± SE), low jitter (5.7 ± 0.1 ms) and high reliability (81.7 ± 2.6%). Here we compared BFCN_BURST_ and BFCN_REG_ and found that both types showed strong response to air puff punishment (Fig. 3c-d). Contrary to previous hypotheses, BFCN_REG_ were also capable of surprisingly fast and precise phasic firing, emitting a precisely timed single action potential, typically followed by a pause and then a reset of their intrinsic theta oscillation (Fig.3c; Fig.S4a). This clearly distinguished them from tonically active striatal interneurons, which did not show such responses (Fig.S4b-d).

BFCN_BURST_ are capable of emitting both bursts of action potentials and single spikes. Therefore, we wondered whether bursts and single spikes represent salient events such as air puffs differently, in which case this should be reflected in a difference in peri-event time histograms of bursts vs. single action potentials aligned to punishment events. We found that bursts of BFCN_BURST_ significantly concentrated after punishment compared to single spikes in most neurons (Fig.3e; Fig.S4e). We observed similar concentration of bursts after reward but not cue stimuli or trial start signals (Fig.S4f-g), suggesting that bursts represent external events differently compared to single spikes.

*In vitro* studies also predicted that tonically active neurons would be more important in controlling slow tonic changes in acetylcholine levels, which could potentially be reflected in higher baseline firing rates of BFCN_REG_. However, we found that baseline firing rates were largely similar across cholinergic cell types and firing patters (median ± SE, BFCN_BURST_-SB, 4.55 ± 1.26; BFCN_BURST_-PL, 5.74 ± 1.39; BFCN_REG_, 3.96 ± 1.0), with slightly faster firing in BFCN_BURST_-PL, consistent with more depolarized membrane potentials and stronger excitatory inputs suggested by our *in vitro* recordings in Fig.2l-o. (BFCN_BURST_-SB vs. BFCN_BURST_-PL, p = 0.11; BFCN_BURST_-SB vs. BFCN_REG_, 0.41; BFCN_BURST_-PL vs. BFCN_REG_, p = 0.0236; Fig. 3f).

### Bursting cholinergic neurons show synchronous activity

Bursts of cholinergic neurons were found to precisely align to reinforcement (Fig.3c-e), generating a strong synchronous activation of the cholinergic system after reward and punishment. Is synchronous firing specific to these unique behaviorally relevant events, or do they occur at other times as well? Synchronous versus asynchronous activation of subcortical inputs have fundamentally different impact on cortical computations. However, while there is a lot known about synchrony in cortical circuits both within and across cell types^26,49–51^, there is little information on synchronous firing in subcortical nuclei. Specifically, no recordings of multiple identified cholinergic neurons have been performed.

In some cases we recorded two (n = 15) or three (n = 3) cholinergic neurons simultaneously, resulting in 24 pairs of concurrent cholinergic recordings. By calculating pairwise cross-correlations we found that BFCN_BURST_, especially BFCN_BURST_-SB, showed strong zero-phase synchrony among each other (6/6 pairs of two BFCN_BURST_-SB and 5/11 pairs containing BFCN_BURST_-SB and PL showed significant co-activation, p < 0.05). BFCN_REG_ showed little synchrony with other BFCN (2/7 pairs that contained at least one BFCN_REG_ were significantly co-activated, p < 0.05; Fig.4a-b, Fig.S5). Co-activation of BFCN_BURST_ typically spanned ±25 ms (27.22 ± 5.37, mean ± SEM; maximum, 42 ms) and was not restricted to the bursts themselves, as single action potentials of bursting neurons showed similar synchrony (Fig.4c); thus BFCN_BURST_ may share a synchronizing input that differentiates them from other BFCN, possibly contributing to the bursting phenotype itself.

**Figure 4.**
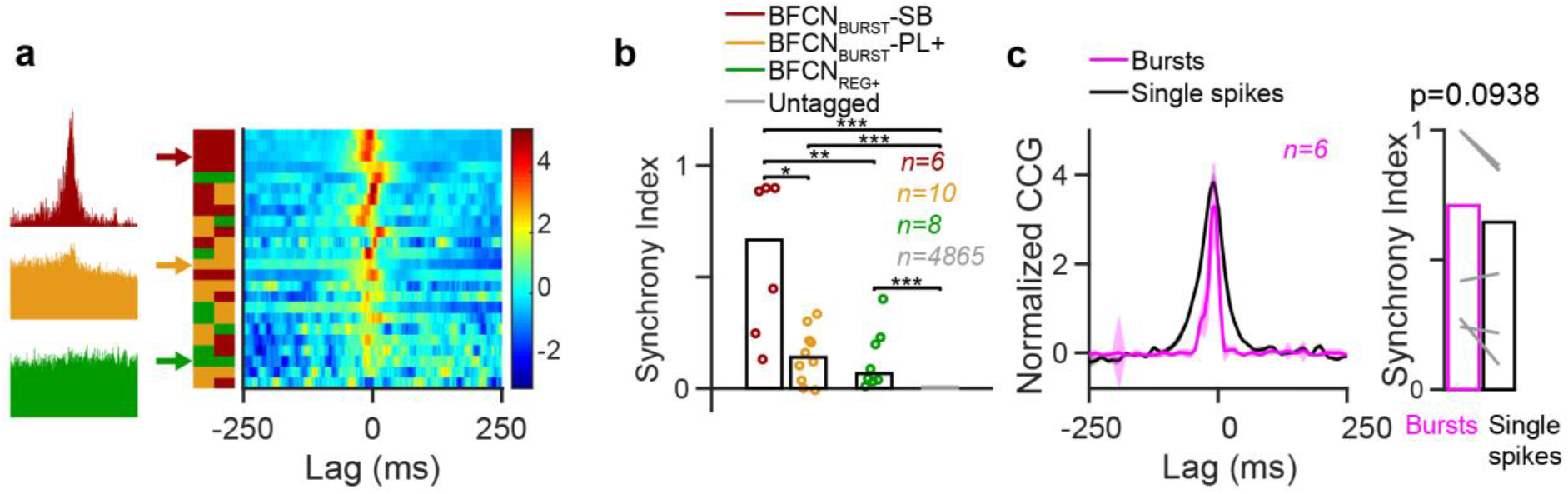
Bursting cholinergic neurons show synchronous activity. **a,** Cross-correlations of pairs of cholinergic neurons. Left, examples. Right, all pairs; left color bar indicates the firing mode of the two neurons that form the pair. Please note, that in some cases the Z-score normalization necessary to show all CCG pairs can magnify central peaks that are otherwise small relative to baseline; therefore, all individual CCG pairs are shown in Fig.S5 without normalization. **b,** Pairs of BFCN_BURST_-SB show stronger synchrony than pairs that contain BFCN_BURST_-PL or BFCN_REG_. Synchrony Index calculated as average cross-correlation in −30-30 ms windows normalized to 100-250 ms baseline period (bars, median). **c,** Both bursts and single spikes of BFCN_BURST_-SB showed zero-lag synchrony.

### Cholinergic bursts are coupled to cortical activity

Cholinergic neurons send dense innervation to the cortex, including projections from the nucleus basalis (NB) to auditory cortices^42,52^. These inputs can potently activate cortical circuits, leading to desycnronization and gamma oscillations^6,53^, which we confirmed by optogenetic stimulation of NB cholinergic neurons that elicited broad band activity in the auditory cortical local field potentials (LFP; Fig.5a). We reasoned that bursts of cholinergic firing might lead to stronger cortical activation, while synchronous activation of ensembles of cholinergic neurons may further increase this effect, providing a finely graded control over cortical activation and thus arousal by the ascending cholinergic system. At the same time, the basal forebrain receives cortical feedback^54–56^ that may be capable of entraining cholinergic neurons thus establishing an ongoing synchrony between cortical and basal forebrain activity, a hypothesis largely under-explored (but see^41,57,58^).

**Figure 5.**
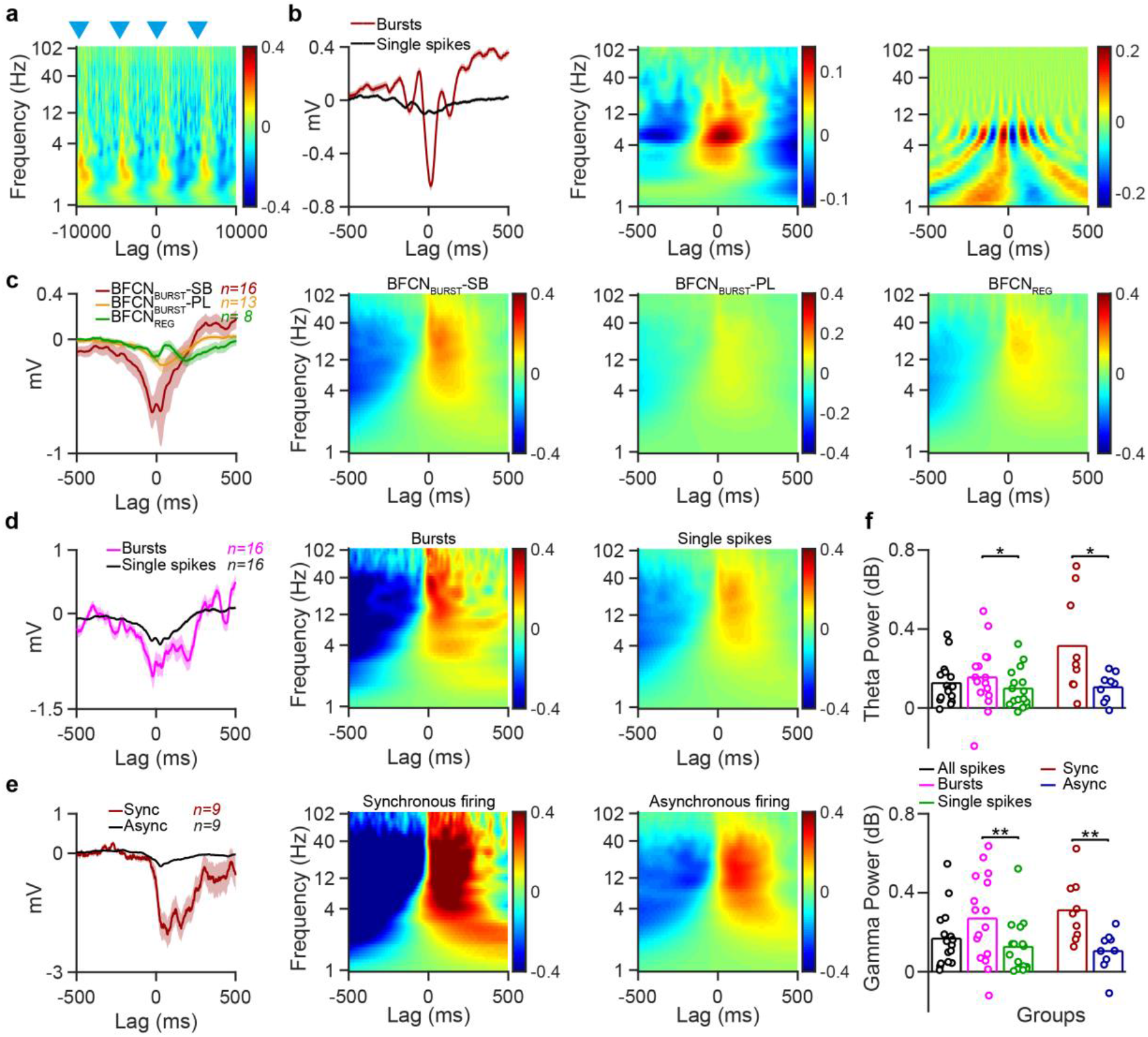
Cholinergic bursts are coupled to cortical activity. **a,** Photostimulation of BFCN activates the auditory cortex; stimulus-triggered average spectrogram aligned to photostimulation (blue triangles). Color code represents spectral power (dB). **b,** Example of a BFCN_BURST_-SB strongly synchronized to cortical theta. Left, STA based on all spikes; middle, STS; right, spike-triggered spectral phase demonstrates phase locking in the theta band. **c,** From left to right: average STA for BFCN groups; average STS for BFCN_BURST_-SB, BFCN_BURST_-PL and BFCN_REG_. **d,** Bursts elicit stronger cortical activation. Left, average STA; middle, average STS for bursts; right, average STS for single spikes. **e,** Synchronous firing elicits stronger cortical activation. Left, average STA; middle, average STS for synchronous firing; right, average STS for asynchronous firing. **f,** Median spectral power in the theta (top) and gamma band (bottom). *, p < 0.05; **, p < 0.01.

To test these possibilities, we calculated spike-triggered LFP averages and spike-triggered spectrogram averages of auditory cortical LFPs aligned to action potentials of BFCN recorded during auditory operant conditioning. We used spike-triggered averages (STA) to identify synchronization between BFCN spiking and cortical oscillations, as LFP changes not phase-locked with BFCN spikes cancel out^41^. Individual STAs aligned to cholinergic spikes showed prominent oscillations in the theta band (4-12 Hz), suggesting that nucleus basalis cholinergic activity can synchronize to cortical theta oscillations (Fig.5b-c). In addition, we often observed strong deflections in cortical LFP after cholinergic spikes (Fig.S6a; peak latency, 36.0 ± 13.0 ms, median ± SE of median) that may be a signature of cortical activation by cholinergic input. To assess this we used spike-triggered spectrograms (STS) to identify evoked responses that are not phase coupled. STS analysis showed high frequency beta/gamma band activity after cholinergic spiking (Fig.5c). Importantly, bursts of BFCN were associated with stronger LFP responses compared to single spikes (Fig.5d,f). We note that a small number of single neurons recorded on the stereotrodes implanted to the auditory cortex showed phase locking to local theta, indicating that oscillations recorded in the auditory cortex were at least partially locally generated (Fig.S7).

Our study confirmed that artificial synchrony of BFCN imposed by optogenetic or electrical stimulation induced cortical desynchronization (Fig.5a), as shown previously^6,53,59^. Since we have found that synchronous activation of BFCN also occurred in a physiological setting (Fig.4), this raises the question whether such synchrony indeed leads to stronger cortical impact. To test this, we focused our analysis on synchronous firing of cholinergic pairs. We found that synchronous events defined by two BFCN_BURST_ firing within 10 ms was associated with strong cortical activation compared to asynchronous firing, confirming our prediction that nucleus basalis signatures of enhanced cholinergic release represent a stronger impact on cortical population activity (Fig.5e-f).

We observed that BFCN_BURST_ often showed synchronization to cortical theta band oscillations (Fig.5b, left). The presence of high values in the theta band in the average spectral phase (phase domain of STS; Fig.5b, right) confirmed this, since it reflects phase-locking to LFP oscillations. We reasoned that differential activation of cholinergic cell types by their inputs might underlie differences in synchronizing with cortical oscillations. It is known that frontal cortical projections to basal forebrain synapse on GABAergic neurons^55,60^, likely providing indirect hyperpolarizing input to cholinergic neurons^4,61^. To model the impact of this circuit on BFCN, we tested whether BFCN_BURST_ and BFCN_REG_ show differential responses to hyperpolarizing current injections *in vitro*. We found that BFCN_BURST_ neurons recovered their spikes with shorter and less variable latency (n = 4, 172.3 ± 9.95 ms, median ± SE of median) than BFCN_REG_ cells (n= 6, 561.25 ± 23.77 ms; p < 0.0001; Fig.S6b-c). This supports the hypothesis that cortically driven indirect inhibition of BFCNs may contribute to their differential coupling to cortical activity.

### Synchrony of BFCN spiking with cortical activity predicts behavior during auditory detection

We have demonstrated that BFCN_BURST_ and BFCN_REG_ are differentially coupled with auditory cortex. However, the functional significance of this connection remains elusive. Therefore, we tested whether synchrony between BFCN and auditory cortex was predictive of behavioral performance during auditory conditioning (Fig.3a). Specifically, we restricted our analysis to one-second-long windows around auditory cue presentation during the operant auditory detection task^24^. We found that BFCN_BURST_, especially BFCN_BURST_-SB, showed larger STA deflections during Hit and False alarm trials compared to Miss and Correct rejection trials (Fig.6a-c). Therefore, synchronization of BFCN_BURST_ with cortical networks predicts mouse responses but not their accuracy, since correct and incorrect responses showed similar STA. In contrast, we found that large STA deflections for BFCN_REG_ specifically predicted Hits; thus, synchronization of BFCN_REG_ and auditory cortex was predictive of performance. We did not find similar predictive activity in a one-second window before the cues, suggesting that predictive synchronization of BF and auditory cortex was evoked by the cue tones. In summary, we found a behavioral dissociation between the two cholinergic cell types; while cortical coupling of BFCN_BURST_ preceded all responses of the animals regardless of performance, BFCN_REG_ specifically predicted correct responses.

**Figure 6.**
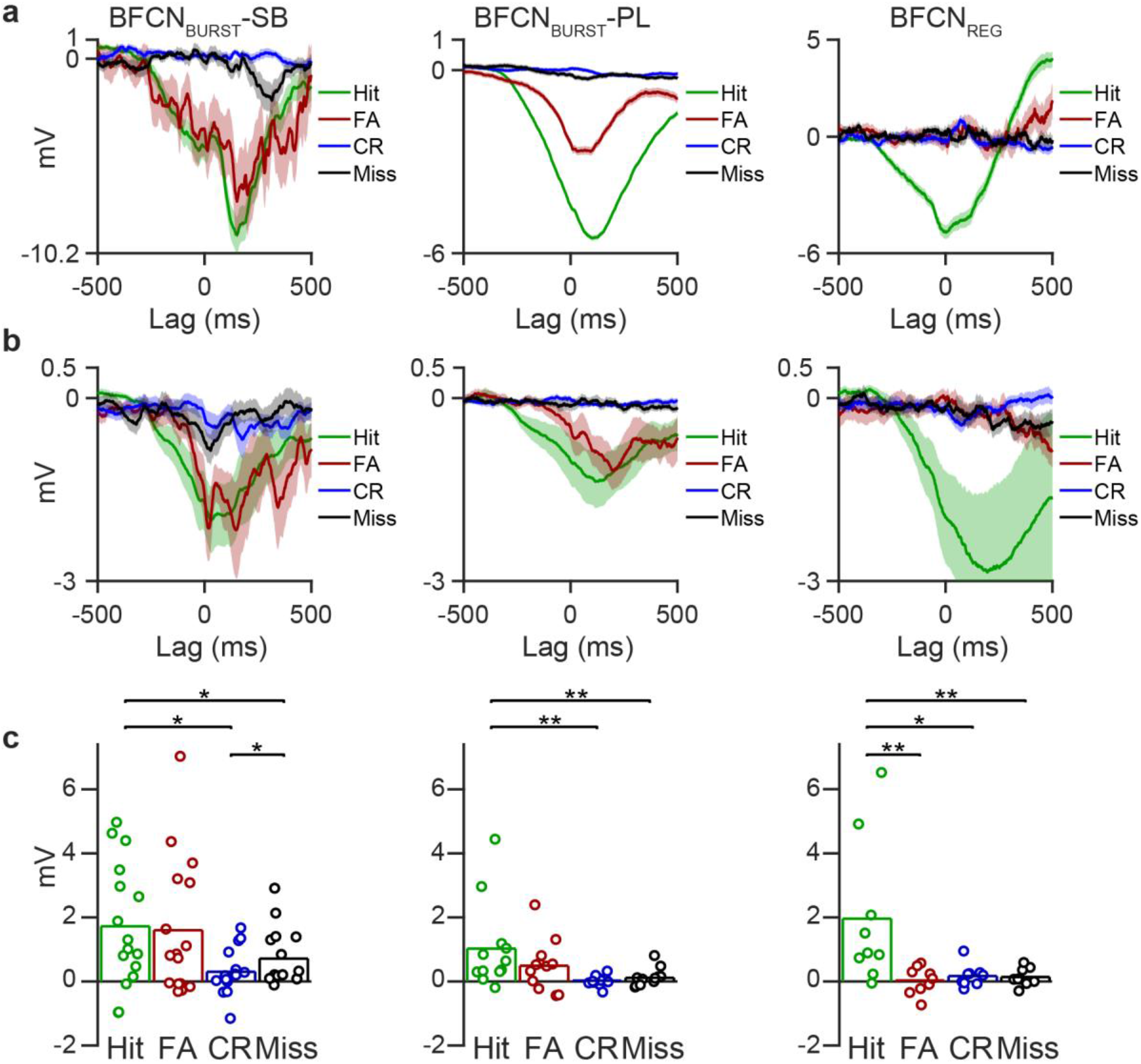
Cortex-BFCN synchrony predicts behavior in an auditory detection task. **a,** Example spike-triggered auditory LFP averages (STA) calculated for spikes of BFCN_BURST_-SB (left), BFCN_BURST_-PL (middle) and BFCN_REG_ (right) restricted to a one-second-long window around cue tone presentations during auditory detection, separated based on trial outcome. FA, false alarm; CR, correct rejection. **b,** Average cue-triggered STA for the BFCN_BURST_-SB (left), BFCN_BURST_-PL (middle) and BFCN_REG_ (right) groups. **c,** Summary comparison of absolute STA deflections for BFCN_BURST_-SB (left), BFCN_BURST_-PL (middle) and BFCN_REG_ (right). *, p < 0.05; **, p < 0.01, Wilcoxon signed rank test.

### The horizontal diagonal band contains few regular cholinergic neurons

We wondered whether the uncovered diversity of cell types is uniform across the basal forebrain; alternatively, differences in the distribution of BFCN_BURST_ and BFCN_REG_ may suggest that dedicated cortical areas are differentially regulated by basal forebrain cholinergic afferents. The cholinergic neurons we recorded were distributed in the nucleus basalis (Fig.1a) and in the more anterior horizontal limb of the diagonal band of Broca (HDB; Fig.7a), spanning nearly 2 mm rostro-caudal distance. This allowed us to investigate whether BFCN types are differentially distributed along the antero-posterior axis of the basal forebrain.

**Figure 7.**
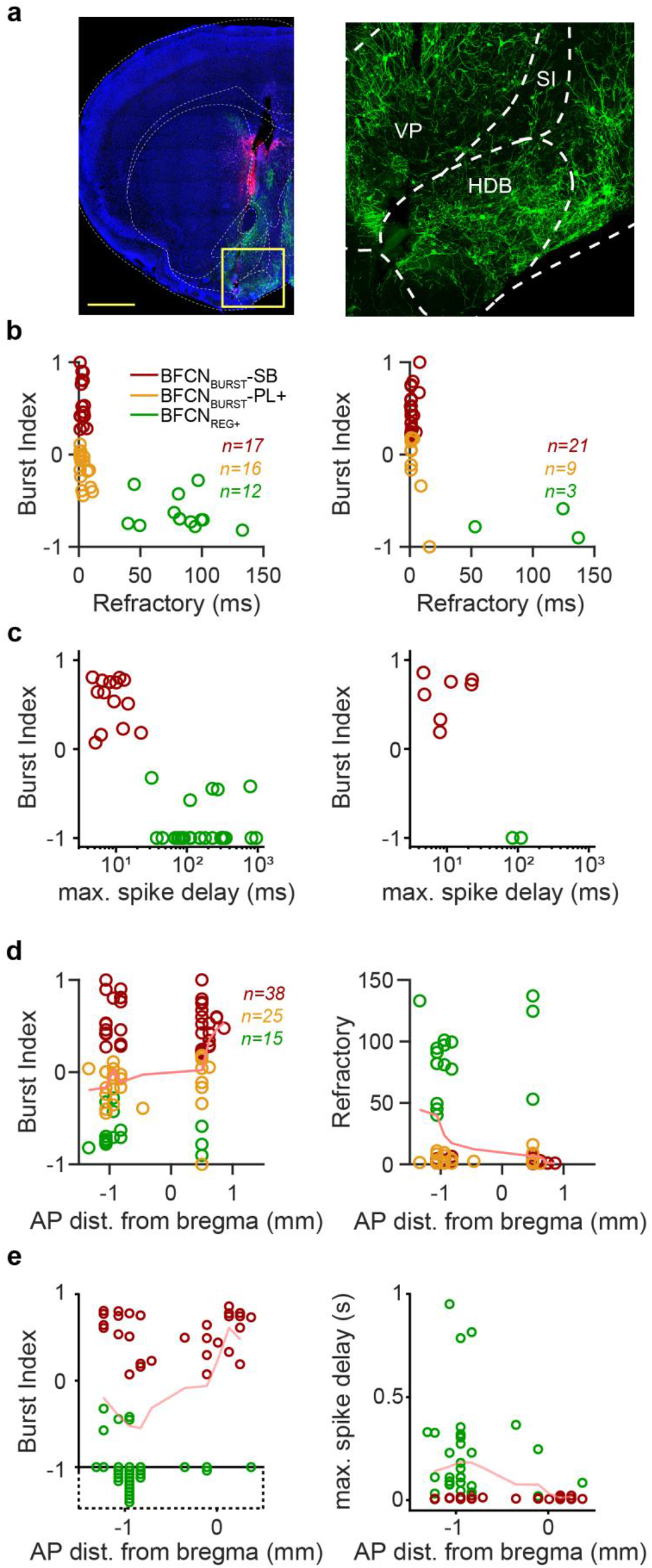
The horizontal diagonal band contains few regular firing cholinergic neurons. **a,** Coronal section showing ChR2 expression (green, eYFP) and tetrode tracks (red, DiI) in the HDB (blue, DAPI staining). Scale bar, 1 mm. Modified from ref. ^24^. **b,** Burst Index vs. relative refractory period for cholinergic neurons recorded from the NB (left) and HDB (right) *in vivo*. **c,** Burst Index vs. maximal spike delay for cholinergic neurons recorded from the NB (left) and HDB (right) *in vitro*. **d,** Burst Index (left) and relative refractory period (right) as a function of antero-posterior localization *in vivo*. **e,** Burst Index (left) and maximal spike delay (right) as a function of antero-posterior localization *in vitro*.

In our *in vivo* recordings, 27% (n = 12/45) of the NB neurons belonged to the regular rhythmic type, while this was only 9% (n = 3/33) for the HDB (Fig.7b). When we recorded NB neurons *in vitro*, 69% (n = 18/26) were BFCN_REG_, whereas only 22% (2/9) BFCN_REG_ was found in the HDB (Fig.7c). The higher proportion of BFCN_BURST_ in our *in vivo* recordings could be due to better cluster separation because of their somewhat higher firing rates (Fig.3f) and distinct spike shape (Fig.2e). Nevertheless, we found that the NB contained three times more regular rhythmic cholinergic neurons both *in vivo* and *in vitro* compared to the HDB that mostly contained the bursting type (p = 0.0007, Chi-square test). In line with these, Burst Index and relative refractory period of cholinergic neurons changed systematically along the antero-posterior axis of the BF (Fig.7d-e), suggesting that different brain areas may receive different combinations of cholinergic inputs.

Turning to untagged HDB neurons we recorded *in vivo*, we found that only 12 out of 560 HDB neurons were characterized as regular firing (Fig.S8), which confirms both the lack of BFCN_REG_ in the HDB (Fig.7) and the connection between regular rhythmic phenotype and cholinergic identity (Fig.1h,Fig.S2h).

## Discussion

We demonstrated that the basal forebrain cholinergic population consist of a burst firing and a regular, rhythmic non-bursting cell type. These types were found both *in vivo* and *in vitro*, and bursts could not be elicited from regular firing cholinergic neurons using a range of current injection protocols. BFCN_BURST_ fired either discrete bursts of action potentials (strongly bursting, BFCN_BURST_-SB) or an irregular pattern of short and long inter-spike intervals resembling a Poisson process (Poisson-like, BFCN_BURST_-PL) depending on their membrane potential and strength of depolarization. Their bursts occurred preferentially after behavioral reinforcement, water reward or air puff punishment, arguing for a separate burst code that selectively represents salient stimuli. BFCN_BURST_ showed strong synchrony among each other and with cortical oscillations, suggesting that they may have a strong impact on cortical processing. Specifically, synchrony between BFCN_BURST_ and auditory cortex at stimulus presentation predicted response timing. In contrast, coupling between BFCN_REG_ and auditory cortex was strongest before mice made successful hits, thus predicting behavioral performance. BFCN_BURST_ and BFCN_REG_ were differentially represented in anterior and posterior basal forebrain. Since anterior and posterior BF have different projection targets^42,62^, distinct brain regions receive different proportions of bursting cholinergic input.

Viewed from the effector side, the cholinergic system plays diverse roles at a variety of temporal scales from slow modulations of sleep-wake cycle^4^ to rapid fluctuations of arousal^6,12,13,63–65^ to instantaneous reactions to salient events serving learning^10,22,24^. This lead to the terminology of ‘tonic’ (seconds to hours) and ‘phasic’ (sub-second) cholinergic effects, demonstrated by amperometric recording of cholinergic signals^12^ and by recording^24^ and imaging^22,23^ cholinergic activity. These findings further inspired the hypothesis that different types of BFCN underlie phasic and tonic effects (Fig.8a). However, another plausible alternative was that different, phasic bursting vs. tonic firing modes of the same neurons are responsible for controlling the time scale of impact (Fig.8b) ^11,13,16^.

**Figure 8.**
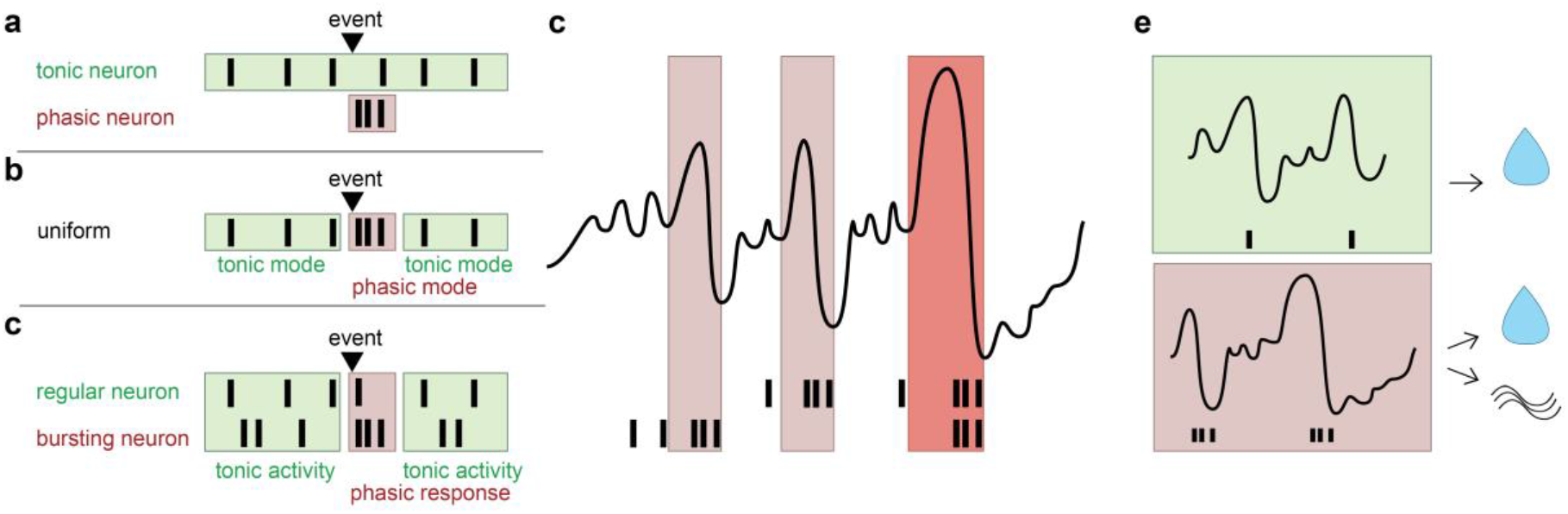
Tonic and phasic cholinergic effects. **a,** Based on heterogeneity found *in vitro*, it was hypothesized that tonic and phasic responses are mediated by different cell types. **b,** Based on homogeneity found *in vivo*, it was suggested that different firing modes of a uniform cell type mediates tonic and phasic effects. **c,** We found that phasic responses to behaviorally significant events are mediated by phasic single spike and burst firing of BFCN_REG_ and BFCN_BURST_, respectively. **d,** Bursts of BFCN synchronize with cortical LFP. Synchronous bursting of BFCN_BURST_ is characterized by stronger synchrony. **e,** Synchronization of BFCN_REG_ to cortical LFP predicts correct detections. Cortical synchronization of BFCN_BURST_ precedes both correct and incorrect responses.

Our result suggests a third, more complex scenario underlying tonic and phasic cholinergic effects (Fig.8c). We propose that there are two basal forebrain cholinergic cell types, demonstrated by our *in vivo* and *in vitro* recordings showing clear separation of regular rhythmic and bursting BFCN (Fig.1e-i, Fig.2b-k). The same neurons produced Poisson-like or strongly bursting firing patterns depending on their membrane potential and synaptic inputs (Fig.2k-n) *in vitro*, showing that BFCN_BURST_-SB and BFCN_BURST_-PL are different firing modes of the same bursting cell type. This claim was also supported by the fact that Burst Index showed a continuum across Poisson-like and strongly bursting firing patterns (Fig.1i). While BFCN_BURST_ and BFCN_REG_ are two separate cell types (Fig.1-2), the firing mode seems crucial to regulating slow and fast cholinergic modulation (Fig.3). Specifically, single spike firing of both cell types contributes to slow tonic modulation by regular theta-rhythmic (BFCN_REG_) or irregular Poisson-like (BFCN_BURST_-PL) firing. Single spike firing also contributes to fast phasic coding by virtue of surprisingly precise spike timing^24,64^ (Fig.3c). In stark contrast, bursts of BFCN_BURST_ selectively enhance phasic responses to salient events (Fig.3e), suggestive of a distinct ‘burst code’ as predicted by theory^66,67^. Nevertheless, it will be important to examine by what mechanisms these cholinergic responses of different temporal scales influence downstream circuits, including synaptic vs. non-synaptic release^68^ and muscarinergic vs. nicotinergic effects^10,69–71^.

Rhythmic BFCN_REG_ beat asynchronously at different frequencies largely independent of each other and BFCN_BURST_ (Fig.4) under our circumstances. The strength of rhythmic firing was correlated with the length of the relative refractory period. Refractoriness may itself contribute to rhythmic firing by imposing regular inter-spike intervals, which may reflect cell-autonomous mechanisms, while extrinsic factors cannot fully be ruled out^72^. Notably, rhythmically firing BFCN_REG_ may share part of the underlying biophysical mechanisms with striatal cholinergic interneurons^72^ and regular firing dopaminergic neurons^73^. The similar auto-frequencies of BFCN_REG_ in the theta range suggest that they may be capable of theta-rhythmic synchronization in a strongly behavior-dependent manner^41^. This was supported by our finding that strong correlations of BFCN_REG_ and auditory population activities in a specific task phase was predictive of mouse performance (Fig.6), which may in part underlie the connection between elevated cortical ACh levels and correct sensory detections^12^. It also suggests that behavior-dependent synchronization may lead to efficient bottom-up information transfer (Fig.8d-e). Similar to this result, careful analysis of behavior-dependent frequency coupling lead to new insight in the active sensing field by revealing behavior-dependent theta-frequency synchronization among hippocampus, respiratory and whisking circuits^74–77^ and prefrontal cortex, suggesting that such theta-frequency binding might be a rather general mechanism.

Unlike BFCN_REG_, activity of BFCN_BURST_ showed strong synchrony across cholinergic neurons and with auditory cortex that was less specific to mouse behavior, predicting both correct and incorrect responses but not performance (Fig.4-6). This suggests that BFCN_BURST_ might convey fast and efficient, although less specific activation of cortical circuits. It is not yet clear whether this unspecific prediction of animal responses is related to stronger sensory perception, task engagement, arousal or other factors that may influence responsiveness irrespective of accuracy. The Poisson-like firing of BFCN_BURST_ neurons might at least in part be a hallmark of internal processing or external sensory events not controlled in our experiments. Indeed, supported mathematically by the Poisson limit theorem, the aggregation of many independent discrete events sum up to a Poisson process, with strong implications to Poisson-randomness found in spike timing even in primary sensory cortices^78^.

In addition, differential rebound response after hyperpolarizing steps in BFCN_BURST_ and BFCN_REG_ *in vitro* suggests that differences in cell type specific properties participate in the mechanisms of basal forebrain-cerebral cortex synchrony. This is in line with previous studies demonstrating that excitatory cortical feedback targets GABAergic inhibitory neurons in the basal forebrain, arguing for a disynaptic inhibition-triggered rebound mechanism for synchronizing BFCN with cortical activity^55^.

As expected, bursts of BFCN_BURST_ were followed by stronger desynchronizations in cortex and predicted an elevation of beta-gamma band activity as compared to single spikes. It is tempting to speculate that fast desynchronization after precisely timed cholinergic bursts might be mediated by fast nicotinic receptors, while muscarinic receptors are more tuned towards slower (‘tonic’) changes of cholinergic levels^10,11,61,79,80^. Within the nicotinic acetylcholine family, α7 receptors may be best suited to mediate fast, precise effects due to their fast kinetics, low open probability and fast recovery^70,81,82^.

The strongest desynchronization was observed after synchronous firing of cholinergic neurons, also indicating that synchrony detected in our paired recordings was likely part of a larger scale synchrony of an ensemble of cholinergic neurons^58,83^. This finding also strengthens a long line of research^6,53,59^ suggesting that synchronous activation of cholinergic neurons leads to strong activation of cortical networks. While previous studies imposed artificial synchrony on the cholinergic system by electrical or optogenetic stimulation, we showed that synchronous cholinergic firing occurs physiologically and this physiological co-firing is indeed associated with a strong cortical impact.

There have been only a handful of *in vivo* recordings of identified BF cholinergic neurons and a consensus view has not emerged. In a seminal juxtacellular labeling experiment, Lee and colleagues recorded cholinergic neurons (n = 5) from the magnocellular preoptic nucleus and substantia innominata and found that cholinergic neurons fire bursts and can synchronize with theta oscillations in the retrosplenial cortex in head-fixed rats^41^. In contrast, Simon et al. labelled cholinergic neurons (n = 3) in the medial septum of anesthetized rats but found different, slow firing patterns without bursts or any correlation with hippocampal theta oscillations^39^. Similarly, Duque et al. recorded cholinergic neurons (n = 3) from substantia innominata and nucleus basalis in anesthetized rats and found slow firing with no synchronization to frontal EEG, n = 1/3 BFCN bursting^40^. In novel study, Guo et al. elegantly demonstrated the learning-related activity of optogenetically identified NB cholinergic neurons, while the firing pattern of optotagged units was not specifically analyzed^84^. Using a larger *in vivo* (n = 78) and *in vitro* (n = 60) data set, we revealed here that these seemingly contradictory results can be reconciled by the presence of two distinct types of cholinergic neurons in the basal forebrain, in line with an earlier *in vitro* study^16^. BFCN_BURST_ show strong synchrony with cortical theta oscillations, whereas synchrony is more behavior-specific and therefore less apparent for BFCN_REG_ despite their intrinsic theta-rhythmic firing.

The BF cholinergic system shows a roughly topographic projection to the cortex and other structures^42,54,56,62,85^. The HDB (also known as the Ch3 group) projects to the olfactory bulb, lateral hypothalamus, piriform cortex, entorhinal cortex^42^ and prefrontal cortices^62^. In contrast, the NB (part of the Ch4 group, which also includes substantia innominata, sometimes included in the NB) projects to the basolateral amygdala and large parts of the neocortex^42,62^. In particular, it strongly innervates lateral parts of the neocortex such as auditory^84,86^, somatosensory and motor cortices, whereas the visual cortex receives its cholinergic innervation from more rostral parts of the basal forebrain^62^. We found BFCN_REG_ to constitute about half (one third to two thirds) of BFCN in the NB, while the more anterior HDB cholinergic neurons were mostly (80-90%) of the BFCN_BURST_ type (Fig.7), suggesting an anatomical difference along the antero-posterior axis of the basal forebrain. Together with our previous paper demonstrating a gradient of valence coding along the dorso-ventral dimension of the nucleus basalis^24^, we uncovered a prominent functional topography of the BF cholinergic system. Added to the large literature of topographical anatomical projections between the basal forebrain and the cortex^42,52,54,56,85,87^, this suggests that basal forebrain inputs, while largely homogeneous with regard to the events they represent, broadcast different messages to their targets in terms of activation strength, synchrony and behavioral function.

Based on theoretical considerations it has been suggested that bursts of spikes may represent distinct stimulus features compared to single action potentials^66,67^, proposing the existence of a separate burst code. Such burst codes have been demonstrated in precise place coding of pyramidal cell complex spikes in CA1, sharp tuning of bursts in visual cortex and visual thalamus or specific coding of complex spikes in Purkinje cells^88–90^. A common theme in these studies is the stronger selectivity, and thus higher signal-to-noise ratio of encoding by bursts vs single spikes. We strengthen this line of research by showing stronger selectivity to salient events by bursts of BFCN, suggesting that the above mechanisms and principles generalize to subcortical networks as well. In addition, Kepecs et al. also predicted that bursts readily synchronize to oscillatory inputs owing to slowly inactivating potassium currents that remain elevated after burst firing^67^. This could serve as a biophysical basis for the stronger synchronization of bursts vs. single spikes to cortical theta oscillations.

## Methods

### Animals

Adult (over 2 months old) ChAT-Cre (N = 15, 14/15 male, Higley et al., 2011), ChAT-ChR2 (N = 3, 3/3 male, Zhao et al., 2011) and PV-Cre (n = 4, 4/4 male) mice were used for behavioral recording experiments under the protocol approved by Cold Spring Harbor Laboratory Institutional Animal Care and Use Committee in accordance with National Institutes of Health regulations. N = 3 male ChAT-Cre mice (over 2 months old) were used for *in vivo* and N = 12 ChAT-Cre (7/12 males, P50-150) mice were used for *in vitro* recordings according to the regulations of the European Community’s Council Directive of November 24, 1986 (86/609/EEC); experimental procedures were reviewed and approved by the Animal Welfare Committee of the Institute of Experimental Medicine, Budapest and by the Committee for Scientific Ethics of Animal Research of the National Food Chain Safety Office.

### *In vivo* electrophysiology and optogenetic tagging experiments

Surgical procedures, viral injection, microdrive construction and implantation, recording, optogenetic tagging and histology were described previously^24^. Mice were trained on one of two versions of an auditory head-fixed detection task. In the operant version, mice had to detect pure tones in a go/no-go paradigm as described in ref. ^24^. In the Pavlovian version, mice responded to reward- and punishment-predicting pure tones with anticipatory licking. In this version, air puff punishment was delivered in a fixed proportion of trials in each trial type, irrespective of the anticipatory lick response of mice^91^.

### Analysis of *in vivo* experiments

Data analysis was performed by built-in and custom written Matlab code (Mathworks) available at https://github.com/hangyabalazs/nb_sync_submitted. The datasets analysed during the current study are available from the corresponding author on reasonable request.

Spike sorting was carried out using MClust (A.D. Redish). Only neurons with isolation distance > 20 and L-ratio < 0.15 were included. Optogenetic tagging was verified by the SALT test. Putative cholinergic neurons were selected based on hierarchical cluster analysis of punishment response properties (response magnitude, PETH correlation with identified cholinergic neurons and PETH similarity scores with templates derived from groups of all unidentified cells and unidentified cells suppressed after punishment). These analyses were described in details previously^24^.

*Auto-correlations* (ACG) were calculated at 0.5 ms resolution. ACG graphs were smoothed by a 5-point (2.5 ms) moving average for plotting. When plotting all or average ACGs per group, individual ACGs were mean-normalized and sorted by Burst Index (bursting BFCN_BURST_) or Refractory (BFCN_REG_). *Burst Index* was calculated following the algorithm introduced by the Buzsaki lab^45^: the difference between maximum ACG for lags 0-10 ms and mean ACG for lags 180-200 ms was normalized by the greater of the two numbers, yielding and index between −1 and 1. *Theta Index* was calculated as the normalized difference between mean ACG for a +-25 ms window around the peak between lags 100 and 200 ms (corresponding to 5-10 Hz theta band) and the mean ACG for lags 225-275 and 65-85 ms. Normalization was performed similarly as for the Burst Index. *Relative refractory period* was defined as low spiking probability after an action potential was fired and calculated by estimating the central gap in the ACG^45^. To estimate the range of delays after an action potential at which spiking happened with lower probability, we calculated the maximal bin count of the ACG smoothed by a 10 ms moving average, and took the delay value at which the smoothed ACG first reached half of this value (width at half height). We note that this definition captures low spike probability and not biophysical partial repolarization, as also used by Royer et al^45^. Since this algorithm allows action potentials in the ‘refractory period’, we used the term ‘relative refractory period’ (lower probability of firing). Nevertheless, this property captured the distinction between regular rhythmic and bursting neurons well (Fig.1). *Cross-correlations* (CCG) were calculated at 1 ms resolution. Segments (±100 ms) after reinforcement events were excluded to avoid trivial event-driven correlations. 0-ms lag (middle) values were excluded to avoid potential contamination from spike sorting artefacts. When plotting all or average CCGs, individual CCGs were Z-scored and smoothed by 15-point moving average. Co-activation was considered significant if raw CCG crossed 95% confidence limits calculated by the shift predictor method for at least two consecutive bins. *Peri-event time histograms* (PETH) were averaged from binned spike rasters and smoothed by a moving average. For comparisons of bursts and single spikes, PETHs were divided by (1 + average baseline PETH). All PETHs were baseline-subtracted for visual comparison.

Local field potential (LFP) recordings were carried out in the primary auditory cortex (A1) simultaneously with the tetrode recordings using platinum-iridium strereotrodes. LFP traces were Z-scored and averaged in windows centered to the action potentials of interest for *Spike Triggered Average* analyses. Positive-deflecting STA traces were inverted before averaging for coherence as depth of recording was not precisely controlled; therefore, we could not draw conclusions from absolute delta phases. Wavelet calculations were performed using Morlet wavelet and *Spike Triggered Spectrograms* were calculated from the wavelet power and phase spectra. Individual frequencies were normalized by their averages to give equal weight to spectral components and visualized on a decibel scale.

Two-sided, non-parametric paired (Wilcoxon signed rank test) or un-paired (Mann-Whitney U-test) hypothesis tests were applied for statistical comparisons as appropriate, unless otherwise stated.

### *In vitro* recordings

Mice were decapitated under deep isoflurane anesthesia. The brain was removed and placed into an ice-cold cutting solution, which had been bubbled with 95% O2–5% CO2 (carbogen gas) for at least 30 min before use. The cutting solution contained the following (in mM): 205 sucrose, 2.5 KCl, 26 NaHCO3, 0.5 CaCl2, 5 MgCl2, 1.25 NaH2PO4, 10 glucose. Coronal slices of 300 μm thickness were cut using a Vibratome (Leica VT1000S). After acute slice preparation, slices were placed into an interface-type holding chamber for recovery. This chamber contained standard ACSF at 35°C that gradually cooled down to room temperature. The ACSF solution contained the following (in mM): 126 NaCl, 2.5 KCl, 26 NaHCO3, 2 CaCl2, 2 MgCl2, 1.25 NaH2PO4, 10 glucose, saturated with 95% O2–5% CO2. Recordings were performed under visual guidance using differential interference contrast (DIC) microscopy (Nikon FN-1) and a 40x water dipping objective. Cholinergic neurons expressing ChR2-mCherry were visualized with the aid of a mercury arc lamp and detected with a CCD camera (Hamamatsu Photonics). Patch pipettes were pulled from borosilicate capillaries (with inner filament, thin walled, OD 1.5) with a PC-10 puller (Narishige). The composition of the intracellular pipette solution was the following (in mM): 110 K-gluconate, 4 NaCl, 20 HEPES, 0.1 EGTA, 10 phosphocreatine, 2 ATP, 0.3 GTP, 3 mg/ml biocytin adjusted to pH 7.3–7.35 using KOH (285– 295 mOsm/L). Recordings were performed with a Multiclamp 700B amplifier (Molecular Devices), low pass filtered at 3 kHz, digitized at 10-20 kHz with NI USB-6353, X Series DAQ, and recorded with an in-house data acquisition and stimulus software (courtesy Attila Gulyás, Institute of Experimental Medicine, Budapest, Hungary). For *in vitro* light illumination, we used a blue laser diode (447 nm, Roithner LaserTechnik GmbH) attached to a single optic fiber (Thorlabs) positioned above the slice.

### Analysis of *in vitro* experiments

All *in vitro* data were processed and analyzed off-line using self-developed programs written in Python 2.7.0 and Delphi 6.0 by A.I.G. and D.S. Spike delay was defined as the time between the start of the one-second-long positive current injection step and the peak time of the first following action potential. Burst frequency was calculated from the following three interspike-intervals. Membrane potential on Fig. 2f-g was calculated as the average membrane potential of a 1-s-long period preceding the positive current injection step. Auto-correlations for each cell were calculated on spikes evoked by step protocols (Fig. 2b) and were smoothed by a 5 ms moving average. In case of Fig. 2n, step protocols form each cell were classified into three groups (Fig. 2n inset). *Burst Indices* were calculated similarly to the *in vivo* recordings: the difference between maximum ACG for lags 0-15 ms and mean ACG for lags 50-300 ms was normalized by the greater of the two numbers, yielding and index between −1 and 1. Average Burst Index as a function of AP distance from bregma was calculated as a 3-section moving average (red line in Fig. 7e).

## Acknowledgement

We thank Drs Janos Szabadics, Viktor Varga, László Acsády, Nóra Hádinger and György Buzsáki for insightful discussions and comments on the manuscript and Katalin Sviatkó for help with graphics in Fig.8. This work was supported by the ‘Lendület’ Program of the Hungarian Academy of Sciences (LP2015-2/2015), NKFIH KH125294 and the European Research Council Starting Grant no. 715043 to BH; NKFIH K115441 and KH124345 to AG; and NINDS RO1NS088661 and McKnight Cognitive Disorders Award to AK; ÚNKP-19-3 New National Excellence Program of the Ministry for Innovation and Technology to PH and EFOP-3.6.3-VEKOP-16-2017-00009 to DS. BH is a member of the FENS-Kavli Network of Excellence.

## Author contributions

BH conceived the project, BH recorded *in vivo* data under the supervision of AK, PH recorded *in vivo* data under the supervision of BH, DS recorded and analyzed *in vitro* data under the supervision of AG and TFF, TL, PH and BH analyzed *in vivo* data, BH, TL and DS wrote the manuscript with comments from all authors.

