## Supplemental material for "Distinct synchronization, cortical coupling and behavioural function of two basal forebrain cholinergic neuron types"

### Supplemental Information

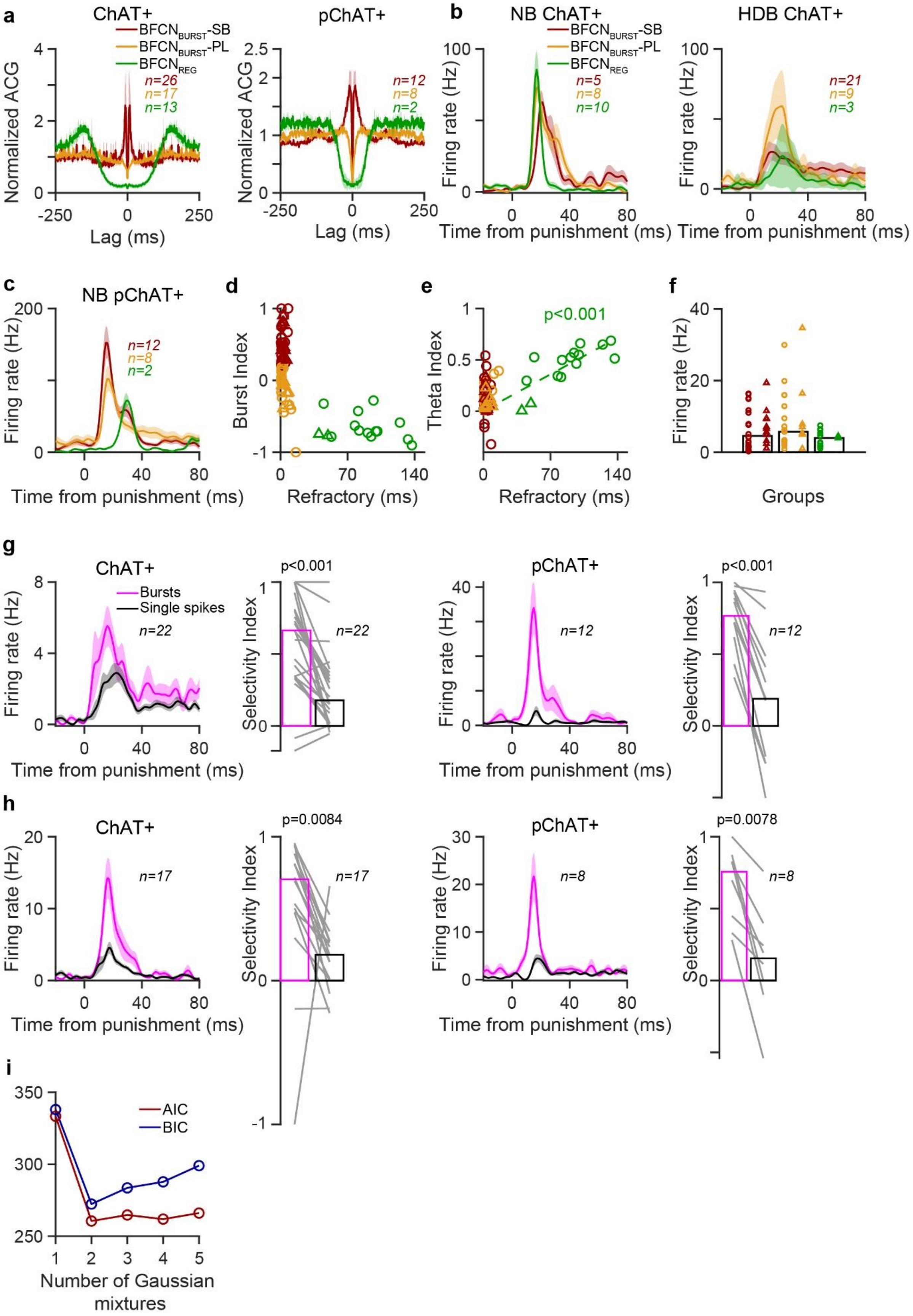
a
ChAT+
BFCN_BURST-SB
BFCN_BURST-PL
BFCN_REG
n=26
n=17
n=13
Normalized ACG
Lag (ms)
pChAT+
n=12
n=8
n=2
b
NB ChAT+
n=5
n=8
n=10
Firing rate (Hz)
Time from punishment (ms)
HDB ChAT+
n=21
n=9
n=3
c
NB pChAT+
d
Burst Index
Refractory (ms)
e
Theta Index
p<0.001
f
Groups
g
Bursts
Single spikes
n=22
Selectivity Index
p<0.001
h
n=17
p=0.0084
p=0.0078
i
AIC
BIC
Number of Gaussian mixtures

**Figure S1. Optogenetically identified and putative cholinergic neurons behave similarly. a,** Average auto-correlogram of $BFCN_{BURST}$-SB (red), $BFCN_{BURST}$-PL (orange) and $BFCN_{REG}$ (green) cholinergic neurons. Left, optogenetically identified; right, putative. While nominal normalized magnitudes may differ due to varying noise levels and moderate sample sizes, the auto-correlation curves are qualitatively similar. **b,** Response to punishment of identified cholinergic neurons (left, identified NB; right, identified HDB). **c,** Response to punishment of putative cholinergic neurons. HDB neurons showed somewhat slower and more variable responses. **d,** Burst Index vs. relative refractory period. **e,** Theta Index vs relative refractory period. No systematic difference between identified (circle) and putative (triangle) cholinergic neurons were detected. **f,** Baseline firing rate did not show systematic differences between identified (circle) and putative (triangle) cholinergic neurons. **g,** Identified (left) and putative (right) $BFCN_{BURST}$-SB exhibited similar burst selectivity. **h,** The same for $BFCN_{BURST}$-PL. **h,** A mixture of Gaussian distributions from 1 to 5 modes were fitted on the logarithm of refractory period distribution. Refractory period of BFCN showed bimodal distribution, confirmed by AIC and BIC model selection measures (lowest value corresponds to best fit model).

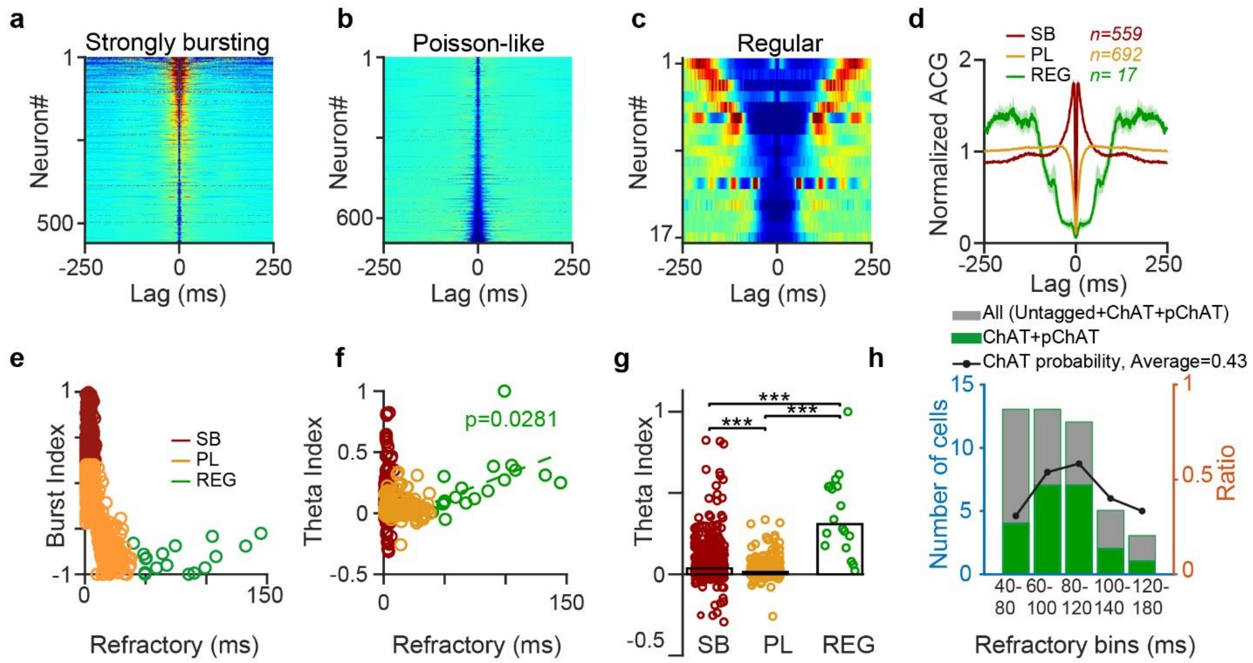


**Figure S2. Regular rhythmic basal forebrain neurons are cholinergic. a-c,** Auto-correlations of untagged bursting (**a**), Poisson-like (**b**), and regular rhythmic (**c**) NB neurons. **d,** Average auto-correlations. **e,** Scatter plot showing Burst Index and refractory period. **f,** Correlation between refractory period and Theta Index ($p = 0.028$ for regular rhythmic neurons). **g,** Median Theta Index. **h,** Predictive value of regular rhythmic firing pattern for cholinergic identity as a function of relative refractory period. Black line and right y-axis correspond to the ratio of (identified or putative) cholinergic neurons to all neurons in the bin. ***, $p < 0.001$.

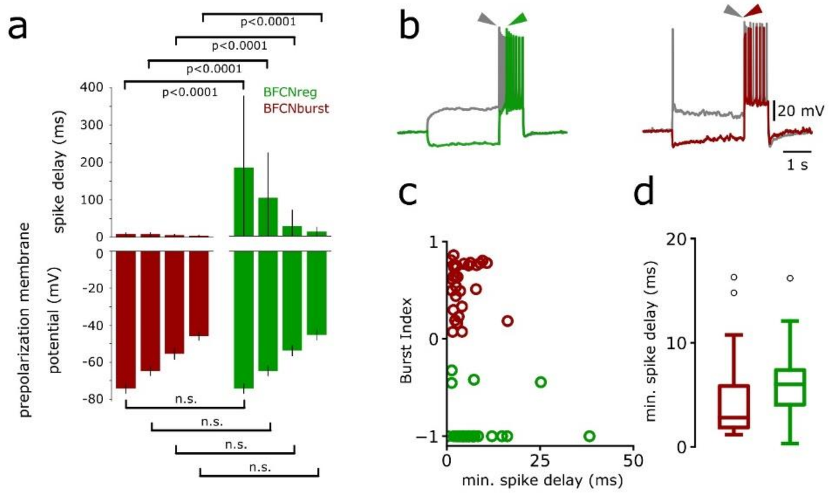


**Figure S3. Similar testing conditions resulted in robust spike delay difference between $BFCN_{BURST}$ and $BFCN_{REG}$ cells, while spike delays were comparable at depolarized membrane potentials. a,** Statistical comparison of spike delay as function of pre-polarization membrane potential. To confirm that late spiking property of $BFCN_{REG}$ was not due to different testing conditions, we compared pre-polarization membrane potentials between groups (two-sample Kolmogorov-Smirnov test; n.s., $p>0.05$). **b,** Example traces of a $BFCN_{REG}$ (left) and $BFCN_{BURST}$ (right) spike response at hyperpolarized and depolarized membrane potentials. Note that the late firing property of $BFCN_{REG}$ is characteristic for hyperpolarized membrane potentials. **c,** Minimum spike delay of each recorded cell vs. Burst Index (green, $BFCN_{REG}$, red; $BFCN_{BURST}$). **d,** Minimum spike delay group statistics.

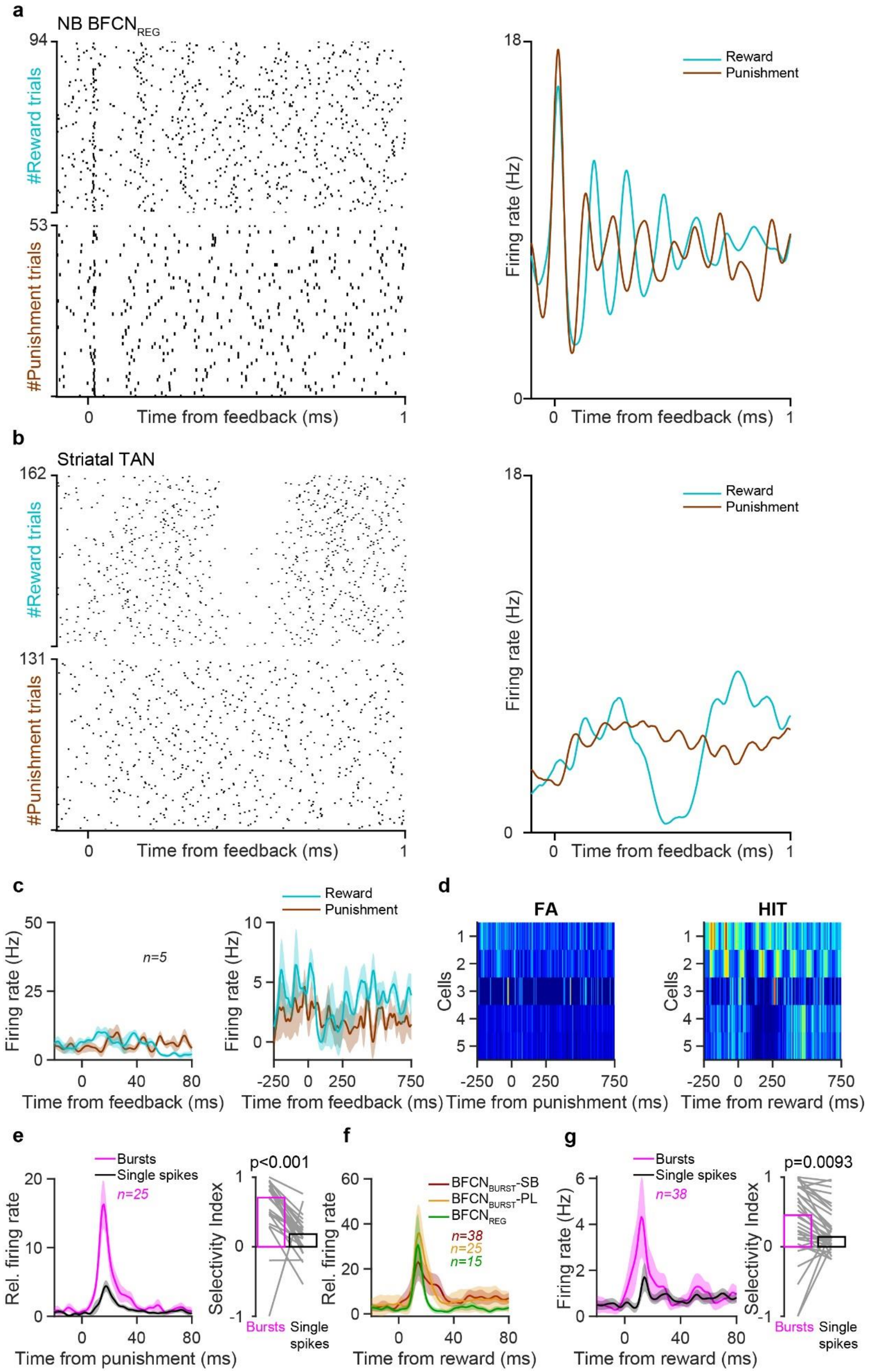
a
NB BFCN$_{REG}$
#Reward trials
#Punishment trials
Time from feedback (ms)
Firing rate (Hz)
Reward
Punishment
b
Striatal TAN
c
n=5
d
FA
HIT
Cells
Time from punishment (ms)
Time from reward (ms)
e
Bursts
Single spikes
n=25
Rel. firing rate
Selectivity Index
p<0.001
f
BFCN$_{BURST}$-SB
BFCN$_{BURST}$-PL
BFCN$_{REG}$
n=38
n=25
n=15
g
n=38
p=0.0093

**Figure S4. Cholinergic bursts transmit phasic information about reinforcers. a,** Raster plots (left) and corresponding peri-event time histograms (PETH, right) aligned to reward and punishment of a $BFCN_{REG}$. After the precise phasic response, the intrinsic theta oscillation resumes. **b,** Raster plots (left) and corresponding PETHs (right) aligned to reward and punishment of an optogenetically identified tonically active cholinergic interneuron (TAN) recorded from the nucleus accumbens. Note the lack of precisely timed action potentials after reinforcement. Instead, TANs show well-characterized so-called 'pause-burst' responses after reward. **c,** Average PETH aligned to reward and punishment at two different time scales of n = 5 optogenetically identified TANs from caudate putamen (n = 3) and nucleus accumbens (n = 2). **d,** PETHs aligned to punishment (left) and reward (right) for all recorder TANs. **e,** $BFCN_{BURST}$-PL cells showed similar burst selectivity after punishment as $BFCN_{BURST}$-SB cells. **f,** BFCN responded phasically to reward. **g,** Bursts of $BFCN_{BURST}$-SB appeared selectively after reward.

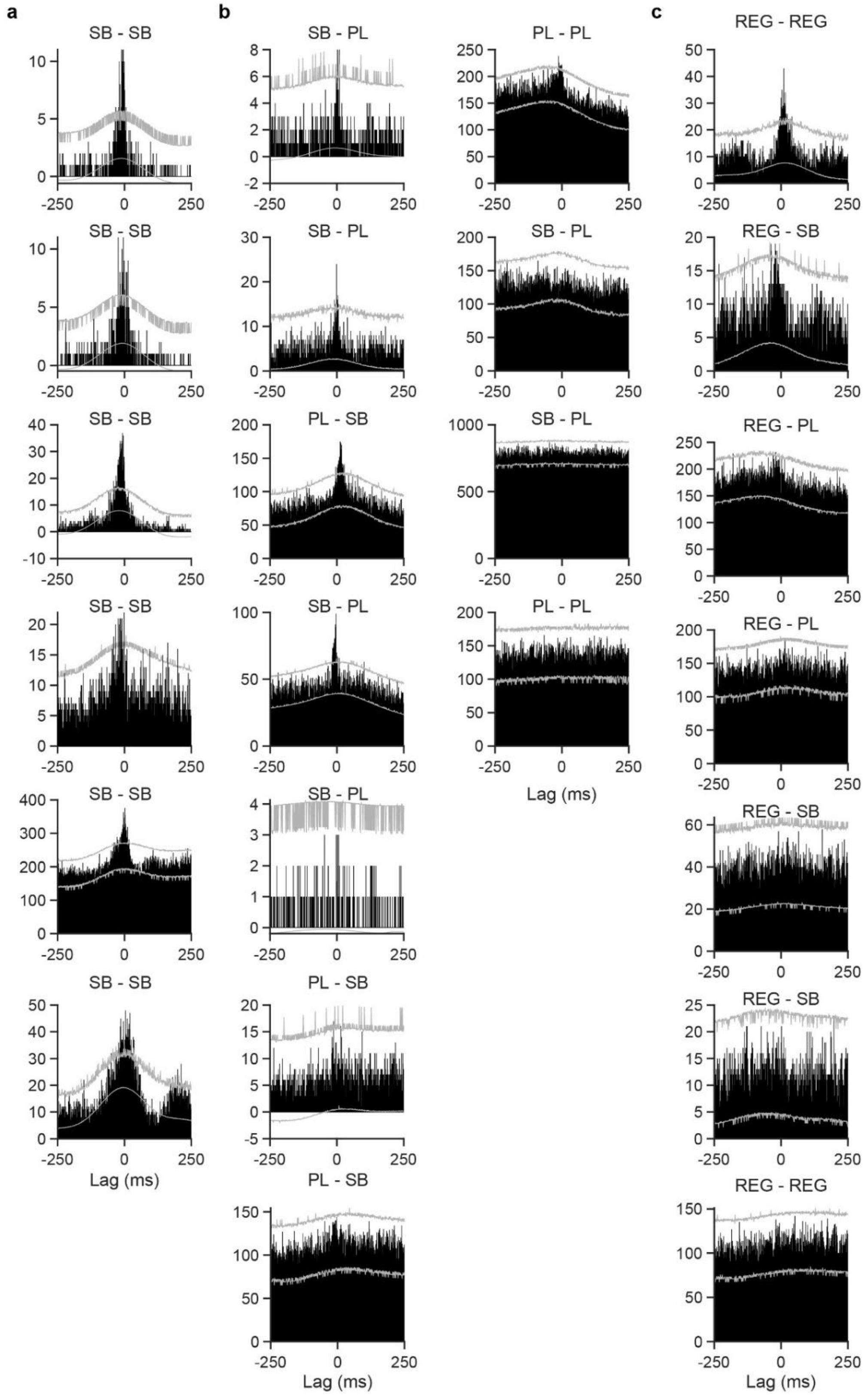


**Figure S5. Individual cross-correlations for all BFCN pairs. a,** Pairs of $BFCN_{BURST}$-SB. **b,** Pairs containing $BFCN_{BURST}$-PL and $BFCN_{BURST}$-SB. **c,** Pairs containing $BFCN_{REG}$.

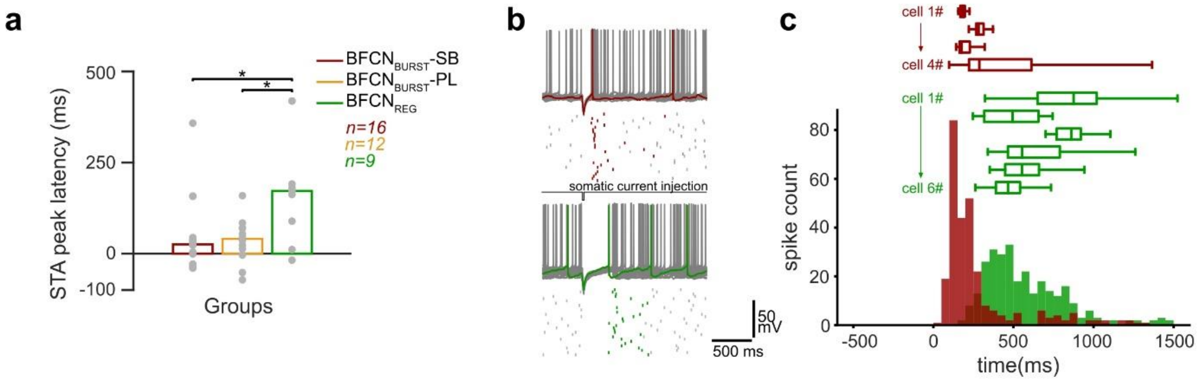


**Figure S6. Bursting and regular rhythmic cholinergic neurons respond differently to hyperpolarization *in vitro*. a,** Peak latency statistics of auditory LFP average triggered on BF spikes *in vivo* (see Fig.5b-c). **b,** Representative responses of a $BFCN_{BURST}$ (top, red) and $BFCN_{REG}$ (bottom, green) upon short (20 ms) hyperpolarizing somatic current injection *in vitro*. Spike rasters of 30 consecutive current injection sessions are displayed below. **c,** Distribution of the first spike latencies following hyperpolarization. Individual cells (horizontal bar plots) are shown above summary histogram (red, $BFCN_{BURST}$; green, $BFCN_{REG}$).

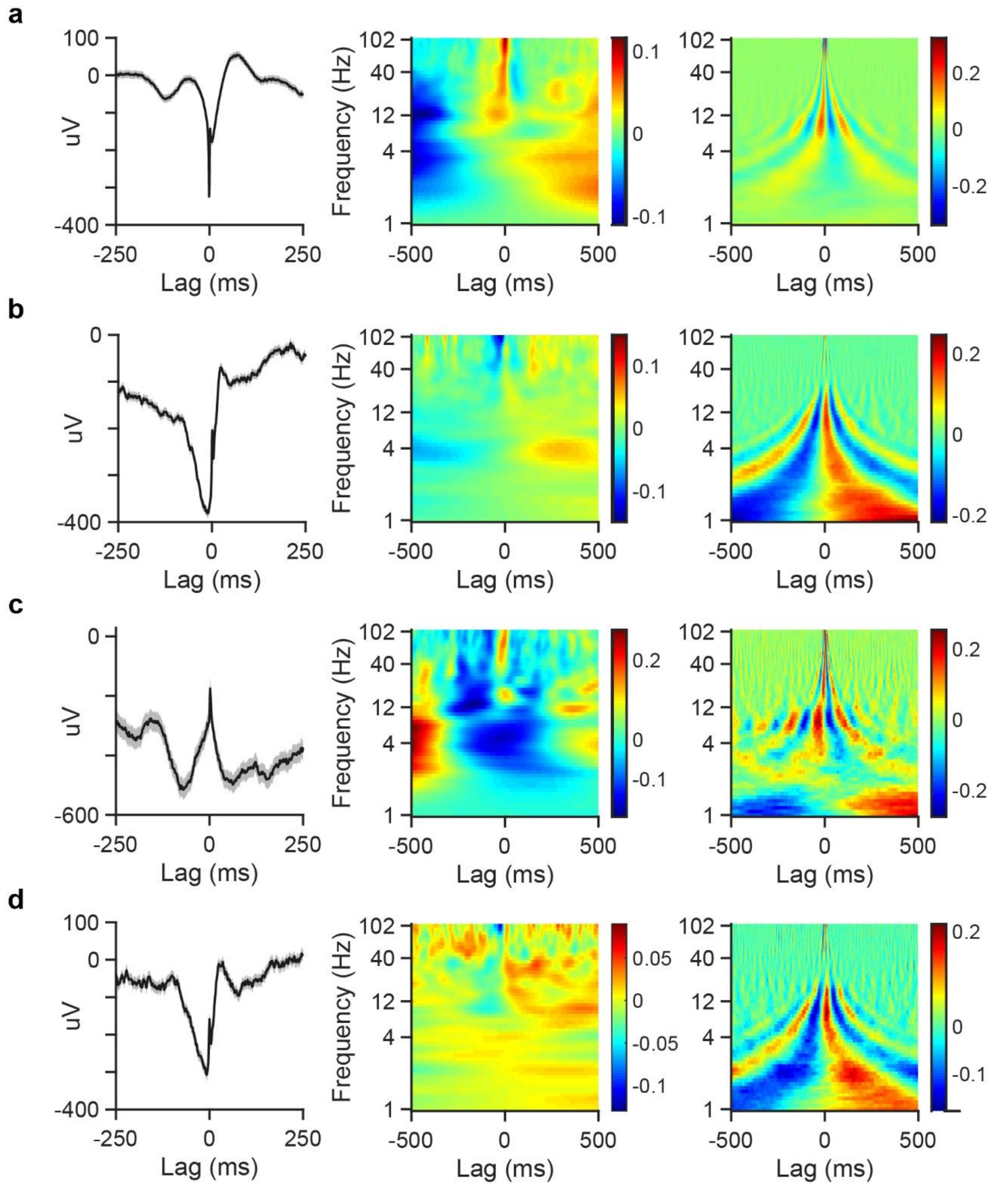


**Figure S7. Some auditory cortical neurons are synchronous with local LFP. a-d,** Example cortical neurons that show synchrony with local LFP. Left, STA; middle, STS power; right, STS phase.

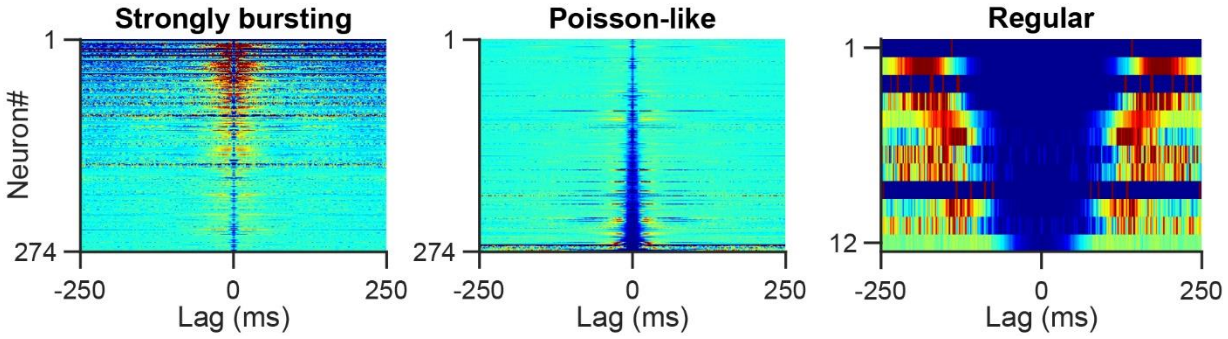


**Figure S8. HDB contains few regular rhythmic neurons.** Auto-correlograms of all unidentified HDB neurons (left, bursting; middle, Poisson-like; right, regular rhythmic). HDB had only 12/560 regular rhythmic neurons.
